# Genomic rearrangements promote diversification of a facultative meiotic parthenogenetic nematode pest (*Meloidogyne graminicola*)

**DOI:** 10.1101/2025.03.20.644268

**Authors:** Ngan Thi Phan, Etienne G.J. Danchin, Georgios D. Koutsovoulos, Corinne Rancurel, Marie-Liesse Vermeire, Guillaume Besnard, Stéphane Bellafiore

## Abstract

*Meloidogyne graminicola* (*Mg*), commonly known as the rice root-knot nematode, is a highly destructive pest that inflicts significant damage on rice crops worldwide. *Mg* is thought to reproduce primarily by meiotic parthenogenesis, but its success across diverse habitats and hosts raises important questions about its adaptation mechanisms, particularly those driving the evolution of its virulence. Documenting the origin of the pathogen, its reproductive strategies and other evolutionary processes shaping its genome are thus crucial to understand its recent, rapid expansion. In this study, we first improved gene annotations of the *Mg* genome to enhance identification of potential secreted parasitism genes. Next, comparative genomics analyses of 13 *Mg* isolates from diverse geographic locations revealed evolutionary changes in the genome, including single nucleotide variations (SNVs), loss of heterozygosity (LoH), and copy number variations (CNVs). These events affected a substantial number of genes, including those coding for secreted proteins, suggesting their roles in nematode adaptation. LoH, the reduction of linkage disequilibrium between SNPs with distance as well as the 4-gamete test all provide evidence for meiotic recombination, supporting some sexual reproduction in *Mg*. Clustering of populations, based on LoH profiles and SNVs, allowed the definition of groups of isolates not correlating with current geographic distribution. The low sequence divergence at the genome level and the lack of clear phylogeographic structure among isolates support the hypothesis of a recent, widespread dissemination of the parasite, especially across Southeast Asia. Overall, our study supports a dual reproductive mode (sexual/asexual) in *Mg*, which offers an evolutionary advantage by balancing clonal proliferation in favorable conditions with occasional sexual reproduction allowing generation of new allelic combinations in adverse environments.

## 1. INTRODUCTION

More than 90% of the rice consumed in the world come from Asia, where around 540 million tons of paddy rice is annually produced [1]. This crop is however increasingly threatened by the prevalence of *Meloidogyne graminicola*, the so-called rice Root-Knot Nematode (RKN). In Asia, RKN is estimated to reduce rice yield by about 15% (reviewed in [2]), but with loss often reaching more than 50%, both in upland and lowland fields [3,4]. These losses may be exacerbated when combined with other biotic or abiotic stresses, such as drought. In addition, the damages caused by this parasite are frequently underestimated due misdiagnosis of the disease or misidentification of abiotic stress, since infected plants may present phenotypes similar to nutritional and water disorders. Not only causing damage in rice, *M. graminicola* is a polyphagous parasite infecting more than 145 host plants, including both cultivated crops and weed species [5]. This devastating parasite is therefore classified as a quarantine pest in several countries (e.g. Brazil, Madagascar, China; [5]) and was added to the EPPO Alert List in Europe in 2017 after being detected in 2016 in northern Italy [6].

Recent surveys show that *M. graminicola* has an almost worldwide distribution. It was first found in the USA [7] and Laos [8] in the sixties, before being identified almost everywhere in South America [9] and Asia [10] where rice is grown, and more recently, in Madagascar [11] and Italy [6]. *Meloidogyne graminicola* has the ability to rapidly increase its population size and spread over a large area in a very short period of time. A freshly hatched juvenile can develop into an adult female laying 250-300 eggs in just 18-21 days under favourable conditions [12–13]. For example, in northern Italy, where this pest has recently been detected, the total infected area has increased by about five folds in just one year (from 19 to 90 ha in 2016-2017; [5]). In Asia, modifications in agricultural practices in response to both climate and socioeconomic changes have also led to a dramatic *M. graminicola* increase [14]. Despite the prevalence of this parasite on a large geographical scale, studies questioning its origin, spread, population structure, and adaptation mechanisms are limited [15–16]. Morphological traits, esterase profiles or molecular data (nuclear ribosomal DNA and mitogenome) have been used to explore the intraspecific variability of *M. graminicola*, but did not reveal any geographical structure among isolates [12, 15–18], suggesting a recent, wide expansion of this parasite. However, these studies based on separate individual markers do not inform on the possible variations at the whole genome level between populations that might be responsible for species adaptation and parasitic success.

Reproduction mode plays a key role in the evolutionary and adaptive potential of parasite species [19]. Sexual reproduction allows combining the genetic material of two parental individuals to produce genetically-diverse offspring. New gene combinations are thus produced, offering opportunities to adapt to changing biotic interaction or environment [20]. Besides, deleterious mutations are more efficiently purged in sexual species via meiotic recombination [21–22]. An animal with clonal reproduction (e.g. parthenogenesis) has poorer adaptability because advantageous alleles from different individuals cannot be combined, and selection efficiency is impaired by the lack of recombination, eventually leading to progressive accumulation of deleterious mutations [23]. Therefore, parthenogenesis was considered an evolutionary dead end. However, asexual reproduction can be advantageous under favorable conditions, and allow a rapid increase of population [24]. In this situation, the genotype adapted to the environment is more easily preserved than under sexual reproduction and recombination. Interestingly, several parthenogenetic species appear to display great adaptability, and comparative genomics has revealed several mechanisms of molecular evolution in their genome [25–26]. The genome structure and sequence of asexual parasites can evolve via movements of transposable elements, loss of heterozygosity, and gene duplications/deletions (i.e. gene copy number variants - CNVs). These genomic variations might be involved in the arms race between the parasite and their hosts [25, 27–29]. For instance, in the mitotic parthenogenetic species *Meloidogyne incognita*, convergent gene loss events have been observed in two isolates that overcame plant resistance [30]. *Meloidogyne graminicola* mainly reproduces through meiotic parthenogenesis. Sexual reproduction has been hypothesized to occur occasionally based on a few evidence of sperm pronuclei inside the eggs (∼0.5%) [31]. Parasites with a mixed reproductive mode are thought to have an evolutionary advantage by combining clonal proliferation, when the genotype is adapted to the environment, and sexual reproduction allowing emergence of new allelic combinations in adverse conditions [32]. If this is verified for *M. graminicola*, it could explain the particularly strong aggressiveness of this pest towards rice in very different agrosystems. However, these hypotheses on the *M. graminicola* reproduction mode are based only on cytogenetic observations [31], and, so far, have never been confirmed at the molecular level. The mechanisms of molecular evolution and their prevalence in the *M. graminicola* genome remain unknown.

In this study, thanks to the recent availability of high-quality *M. graminicola* genome sequences [33, 34, 35], genes, including those coding for secreted proteins, were annotated at a high level of completeness. Comparative genomics of isolates sampled over a large area was conducted to (i) investigate the global genetic diversity at the genome level, and (ii) search for molecular signatures of evolution of the genome. To address these questions, we have sequenced the genomes of 13 isolates mostly distributed in South-East Asia. Genomes of these isolates were compared to the reference genome of isolate Mg-VN18 [33] to detect sequence variants such as single nucleotide variants (SNVs), loss of heterozygosity (LoH), and copy number variants (CNVs). SNVs identified among isolates were used to study the species phylogeography and to test for evidence of recombination. The results of our study revealed that *M. graminicola* populations exhibit evidence for recombination and numerous genomic reorganization with potential impact on their adaptive evolution. This parasite also likely evolved a specific mechanism allowing it to adapt to changing environments through gains or losses of gene copies. The absence of a clear phylogeographic signal based on genomic variants, similarly to previous observations on the mitochondrial genomes, reinforces the hypothesis of a recent spread of this species at least in South East Asia.

## 2. RESULTS

### 2.1 Enhanced genome annotation results in higher completeness and improved secretome prediction

Eugene predicted 15,518 genes spanning 26.5 Mb (63.86%) of the genome of *Meloidogyne graminicola* (*Mg*). They include 15,167 protein-coding genes and 351 non-coding genes (Table 1). Protein-coding genes had an average length of 1,733 bp and covered 14.7 Mb (35.42%) of the genome. Introns were detected in 94% of them with an average of 5.7 introns per gene. The GC percent in coding regions was 30.8% while an even lower GC content (17.4%) was found in introns. Canonical splice donor (GT) and acceptor (AG) sites were identified by Eugene and no non-canonical site was predicted in contrast to the previous annotation done with Maker [33]. Non-canonical splice sites were also not identified in the genomes of other *Meloidogyne* species (i.e. *M. incognita*, *M. javanica*, and *M. arenaria*; [77]), supporting the accuracy of the Eugene annotation for the *Mg* reference genome. The average space between two protein-coding genes was 973 bp. The new gene prediction using Eugene (Eugene_Mg proteome) significantly improved completeness compared to the previously predicted proteome ASM1477313v1 generated with Maker [33] across all BUSCO databases. For instance, with the eukaryote_odb10 database, the C-scores were 69.4% for Eugene_Mg versus 57.3% for ASM1477313v1 (Figure S1). Although Eugene annotated a higher number of proteins (15,518) compared to the published proteomes PRJNA411966 (14,062 proteins; [34]) and T2T (12,968 proteins; [35]), its BUSCO metazoan C-score was 2.1% lower than that of PRJNA411966 and approximately 9.9% lower than the haploid telomere-to-telomere (T2T) genome.

**Table 1.**
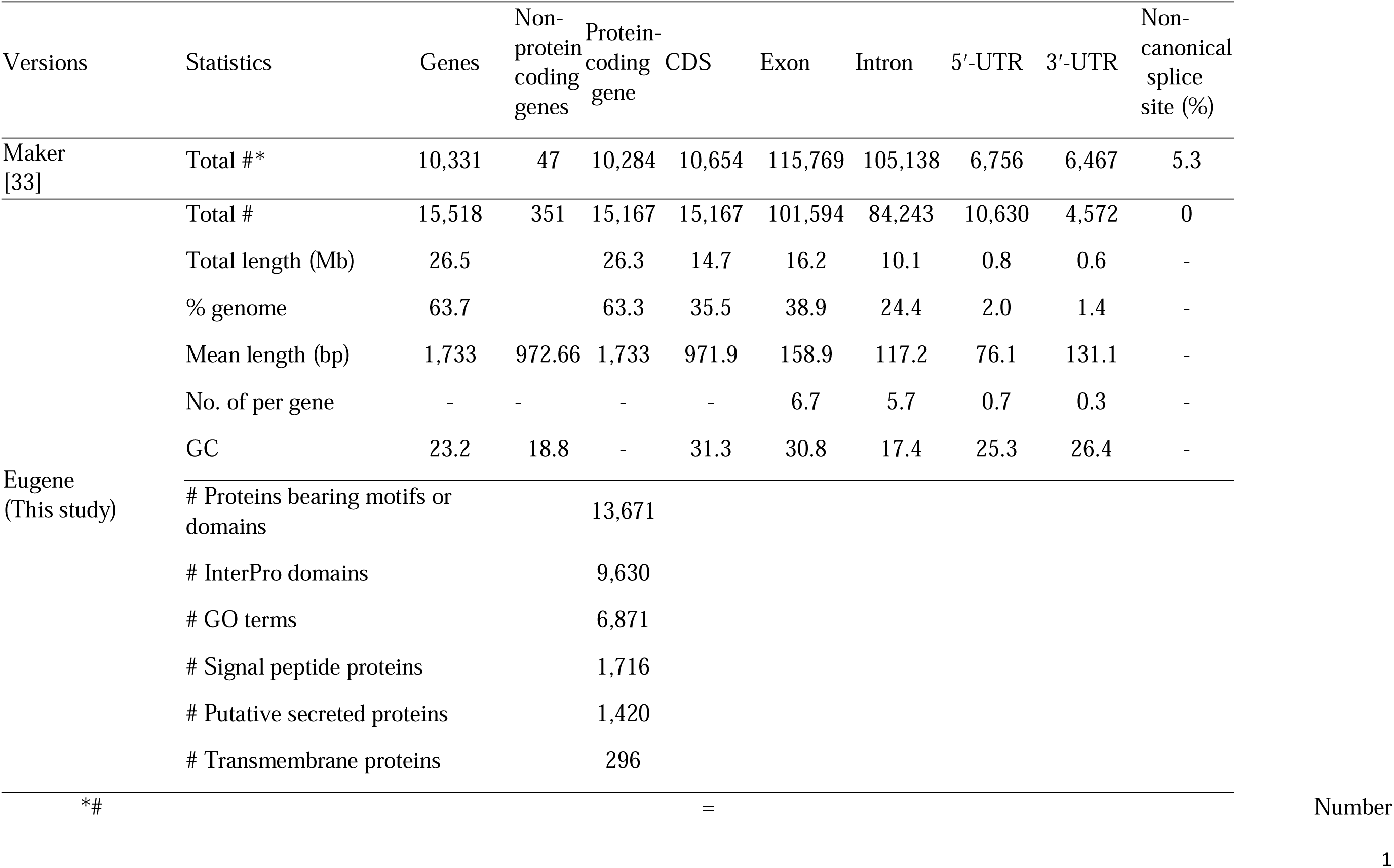
General statistics of gene annotation for the *M. graminicola* genome sequence in this study, and comparison with the previous version annotated by Phan et al. [33].

A total of 1,716 proteins were predicted to bear a signal peptide for secretion, of which 296 also had at least one predicted transmembrane helix and were eliminated, resulting in 1,420 possibly secreted proteins (*PSP*; Table 1). *PSP* are likely to encompass effector proteins secreted by nematodes in plant tissue and thus to play an important role for parasitism [78]. Protein motifs or domains were identified in 13,671 proteins. A total of 7,027 different InterPro domains were identified in 9,630 proteins, allowing further functional annotation with GO terms and metabolic pathways. GO terms were assigned to 6,871 proteins based on the presence of InterPro domains (Table 1).

### 2.2 Contrasting loss of heterozygosity among the 13 *Mg* isolates

Variant calling enabled the detection of 693,013 SNVs among the 13 *Mg* isolates, including both homozygous and heterozygous variant genotypes. Visualizing the positions of these variants on scaffolds revealed variable patterns among isolates (Figures 1 and S2). Indeed, variants are not uniformly distributed but instead enrichments (SNV clusters) are visible in certain genomic regions. In particular, at the same positions, some isolates show ranges of homozygous variants (“1/1” or “0/0”), while other isolates have a heterozygous status (“0/1”). For example, at position 20 to 55 kb in contig mg58 (Figure S2A), Mg-Java2 and Mg-Borneo show a homozygous region with the reference haplotype (“0/0”), Mg-VN27 has a homozygous region with the alternative haplotype (“1/1”), while others were heterozygous (“0/1”). Another clear example for this observation could be also found at the position 100 to 150 kb in contig mg78 (Figure 1). Genomic regions with such a contrasted pattern of rare homozygosity versus frequent heterozygosity among isolates at a given locus most likely result from the fixation of one allelic variant [= loss of heterozygosity (LoH)]. By mapping HiSeq reads of Mg-VN18 against its haploid reference genome as an internal control, we identified stretches of homozygous loci (“0/0”) in Mg-VN18, while other isolates exhibited heterozygous loci (“0/1”), indicating loss of heterozygosity (LoH) in Mg-VN18. For example, from positions 0 to 25 kb in contig mg137, both Mg-VN18 and Mg-C25 were homozygous (0/0), whereas all other isolates were heterozygous on this region (0/1) (Figure S2B). All DNA fragments that have reached fixation were recorded (Table 2), showing that the number of LoH events varied from 7 to 55 among isolates, covering from 0.06% to 7.01% of the genome. The LoH region sizes varied from 1 to 613.13 kb. LoH regions shorter than 10 kb were identified in all isolates, while LoH regions larger than 100 kb were observed in most isolates, except Mg-Brazil, Mg-VN11, and Mg-VN18 (Table 2, Figure S9). Interestingly, these large LoH regions were often located at the ends of the scaffolds—for example, on scaffolds mg24, mg33, and mg47 in the Mg-Bali isolate—suggesting that these regions may correspond to the termini of homologous chromosomes that are often called terminal LoH (Figure S9).

**Figure 1.**
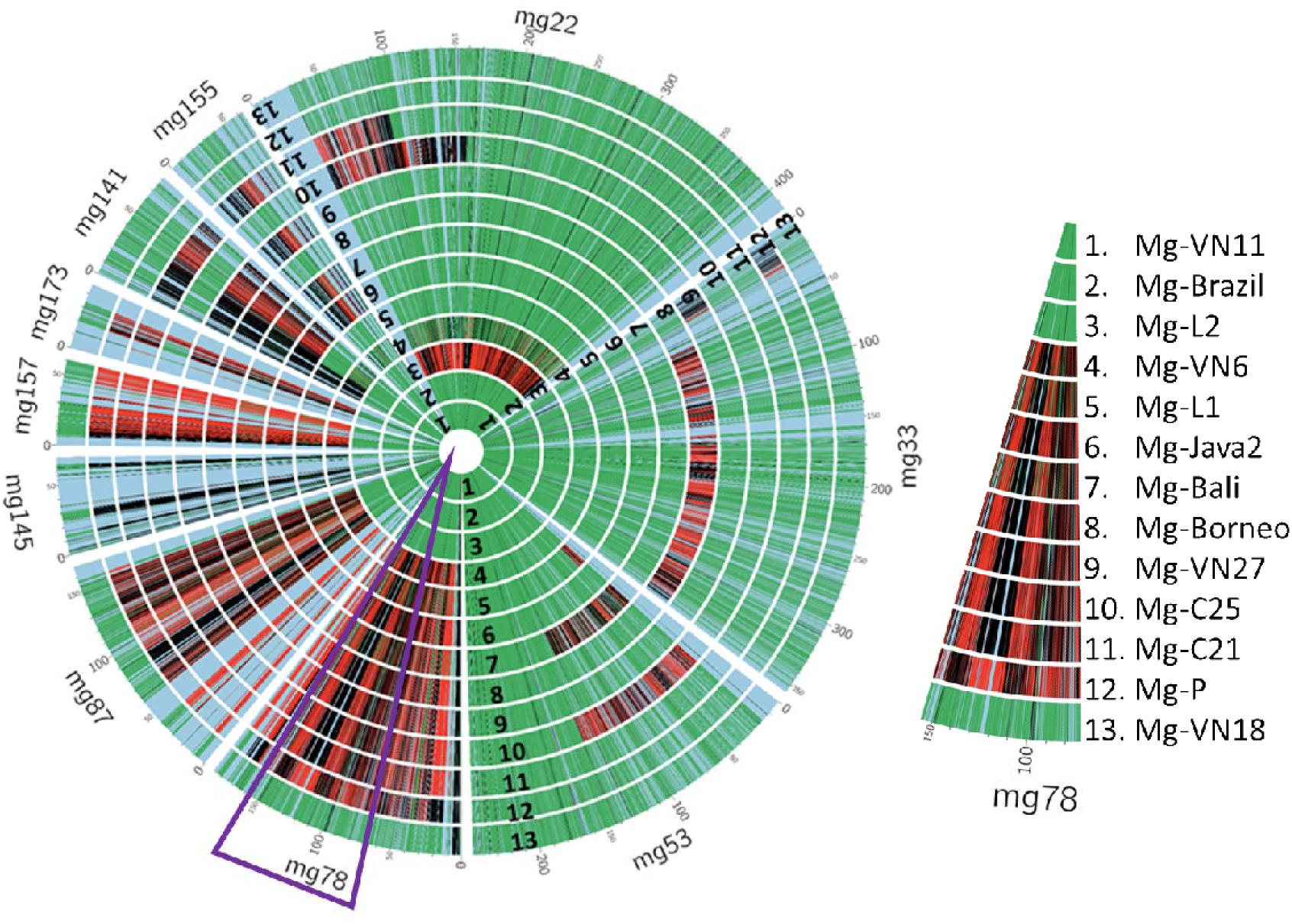
Presentation of loss of heterozygosity (LoH) regions on the 13 *Mg* isolates on ten scaffolds in which LoH were common among isolates. The length of scaffolds is represented in the scale. Each inner circle represents the genome sequence of an isolate (1 to 13). Each radial line indicates the position of SNV on the scaffold. The color of each radial line indicates genotype of SNV with the following color code: green = heterozygous state (genotype “0/1”); black = homozygous state with reference haplotype (genotype “0/0”); red = homozygous state with the alternative haplotype (genotype “1/1”). Therefore, the “green” regions indicate heterozygous state while the “red and black” regions indicate loss of heterozygosity (LoH) regions. The blue regions indicate conserved homozygous (0/0) regions between isolates.

**Table 2.**
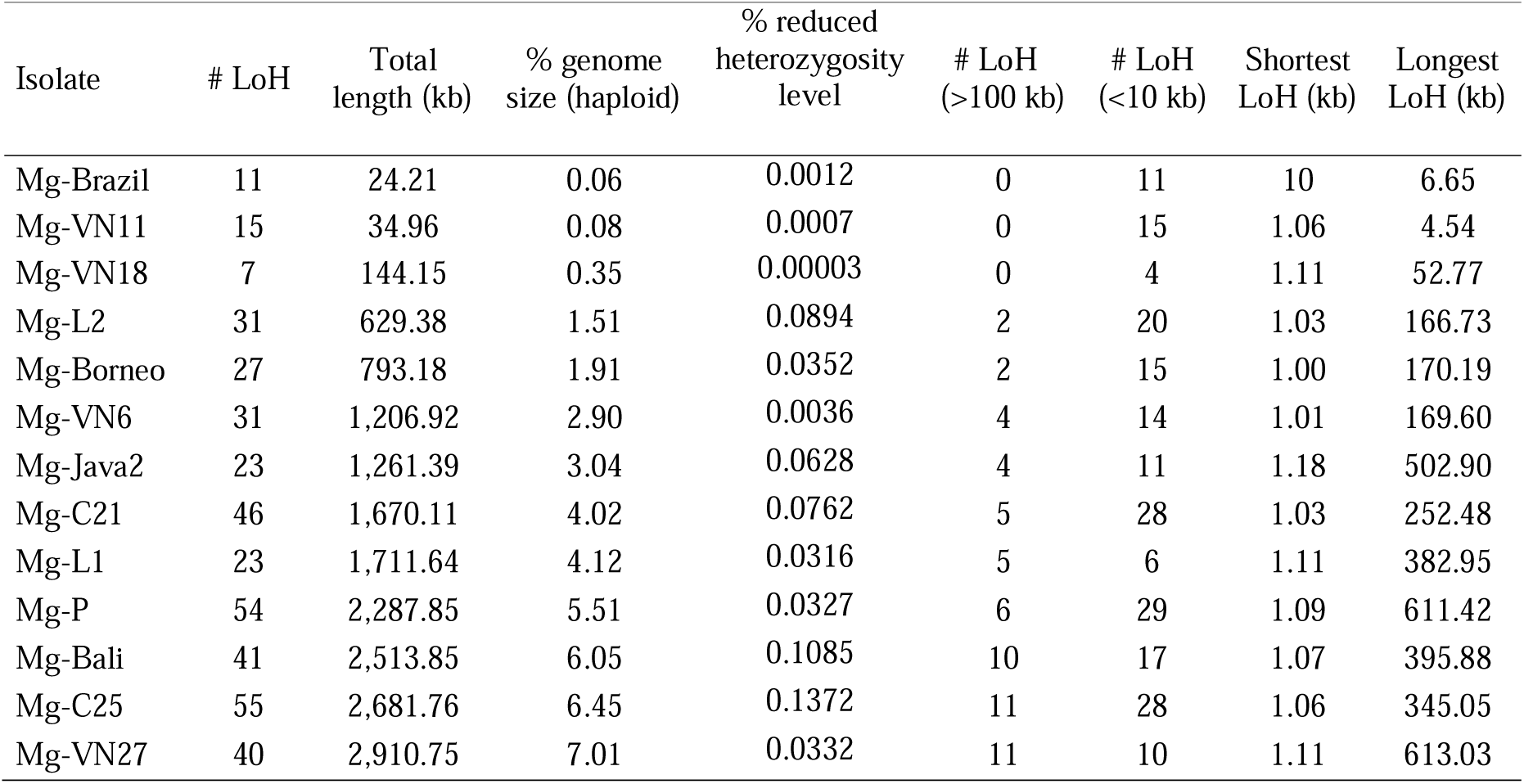
Number and total length of genomic regions that have reached fixation (loss of heterozygosity) among the 13 *M. graminicola* isolates.

| Isolate | # LoH | Total length (kb) | % genome size (haploid) | % reduced heterozygosity level | # LoH (>100 kb) | # LoH (<10 kb) | Shortest LoH (kb) | Longest LoH (kb) |
| --- | --- | --- | --- | --- | --- | --- | --- | --- |
| Mg-Brazil | 11 | 24.21 | 0.06 | 0.0012 | 0 | 11 | 10 | 6.65 |
| Mg-VN11 | 15 | 34.96 | 0.08 | 0.0007 | 0 | 15 | 1.06 | 4.54 |
| Mg-VN18 | 7 | 144.15 | 0.35 | 0.00003 | 0 | 4 | 1.11 | 52.77 |
| Mg-L2 | 31 | 629.38 | 1.51 | 0.0894 | 2 | 20 | 1.03 | 166.73 |
| Mg-Borneo | 27 | 793.18 | 1.91 | 0.0352 | 2 | 15 | 1.00 | 170.19 |
| Mg-VN6 | 31 | 1,206.92 | 2.90 | 0.0036 | 4 | 14 | 1.01 | 169.60 |
| Mg-Java2 | 23 | 1,261.39 | 3.04 | 0.0628 | 4 | 11 | 1.18 | 502.90 |
| Mg-C21 | 46 | 1,670.11 | 4.02 | 0.0762 | 5 | 28 | 1.03 | 252.48 |
| Mg-L1 | 23 | 1,711.64 | 4.12 | 0.0316 | 5 | 6 | 1.11 | 382.95 |
| Mg-P | 54 | 2,287.85 | 5.51 | 0.0327 | 6 | 29 | 1.09 | 611.42 |
| Mg-Bali | 41 | 2,513.85 | 6.05 | 0.1085 | 10 | 17 | 1.07 | 395.88 |
| Mg-C25 | 55 | 2,681.76 | 6.45 | 0.1372 | 11 | 28 | 1.06 | 345.05 |
| Mg-VN27 | 40 | 2,910.75 | 7.01 | 0.0332 | 11 | 10 | 1.11 | 613.03 |

Although the LoH regions accounted for up to 7.01% of the genome size, they caused only a minimal reduction in heterozygosity levels, ranging from 0.00003% (Mg-VN18) to 0.1372% (Mg-C25), in the genome of each isolate (Table 2). The genome heterozygosity level of each isolate, calculated at *k* = 21, ranged from 1.78% (Mg-L2) to 3.40% (Mg-C21; Figure S3). These values remained consistent across different *k*-mer sizes (*k* = 19, 21, 25, and 27; Figure S3).

A dendrogram based on overlapping LoH regions among the 13 isolates revealed two main groups: Group 1 consisted of Mg-VN18, Mg-VN11, Mg-L2, and Mg-Brazil; Group 2 contained nine isolates and was divided into two subgroups: Sub-group 2.1 containing Mg- C25 and Mg-P; and Sub-group 2.2, encompassing the remaining seven isolates (Figure 2). Interestingly, the nine isolates from Group 2 shared LoH events on scaffolds mg78, mg87, mg145, mg157, and mg173 for a total of 490 kb (∼1.18% of the genome size; Figure 1). Among these nine isolates, Mg-P has a distinct haplotype at scaffold mg78 compared to the others (Figure 1). Isolates from the same countries, for example Laos (Mg-L1 and Mg-L2), Cambodia (Mg-C21, Mg-C25), and Vietnam (Mg-VN27, Mg-VN11, Mg-VN6), showed contrasting LoH profiles (Figure 2). Meanwhile in Group 1, isolates from distant continents, Asia and America, shared a similar LoH profile. Therefore, the LoH phenomenon showed no clear association with the geographic origin of isolates (Figures 1 and 2).

**Figure 2.**
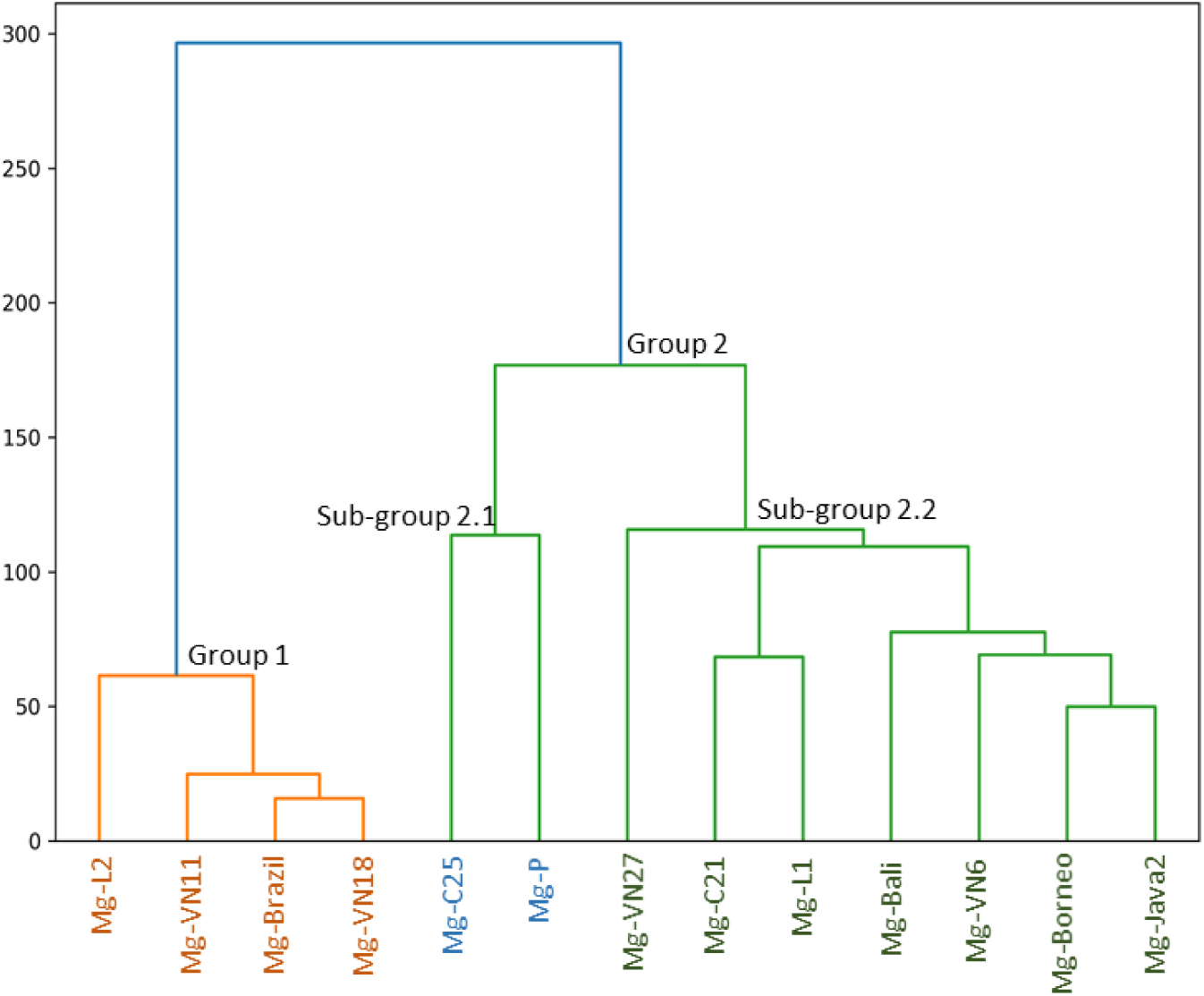
Hierarchical clustering of the 13 *Mg* isolates based on LoH regions.

A total of 3,639 protein-coding genes were potentially affected by LoH, including 381 genes predicted to encode secreted proteins (*PSP* genes; Table 4). Non-synonymous and synonymous SNPs were found in 2,115 protein-coding genes, including 219 *PSP* genes. The proportion of secreted protein-coding genes compared to non-secreted protein-coding genes affected by LoH (#PSP/#nonPSP) did not differ significantly from the proportion in the rest of the genome with values of 11.4% ± 0.05 versus 10.0% ± 0.002, respectively (Figure 3). However, the distribution of ratio of non-synonymous to synonymous substitutions (dN/dS) in *PSP* genes located within LoH regions was different and shifted towards higher values than the distribution observed for *PSP* genes located in the rest of the genome for each isolate (average values 1.40 ± 0.20 versus 1.14 ± 0.04, *P* < 0.05; Figure 3). Additionally, the dN/dS ratio for *PSP* genes in LoH regions was significantly higher than that for non-*PSP* genes in the same regions (1.40 ± 0.20 versus 0.63 ± 0.06, *P* < 0.001; Figure 3). The nucleotide diversity at synonymous sites (π_s_), nonsynonymous sites (π_n_), and the selection efficiency ratio (π_n_/π_s_) were similar between LoH regions and the whole genome. This consistency was observed for both protein-coding genes (π_n_/π_s_ ratio ∼0.20) and secreted protein-coding genes (π_n_/π_s_ ratio ∼0.29).

**Figure 3.**
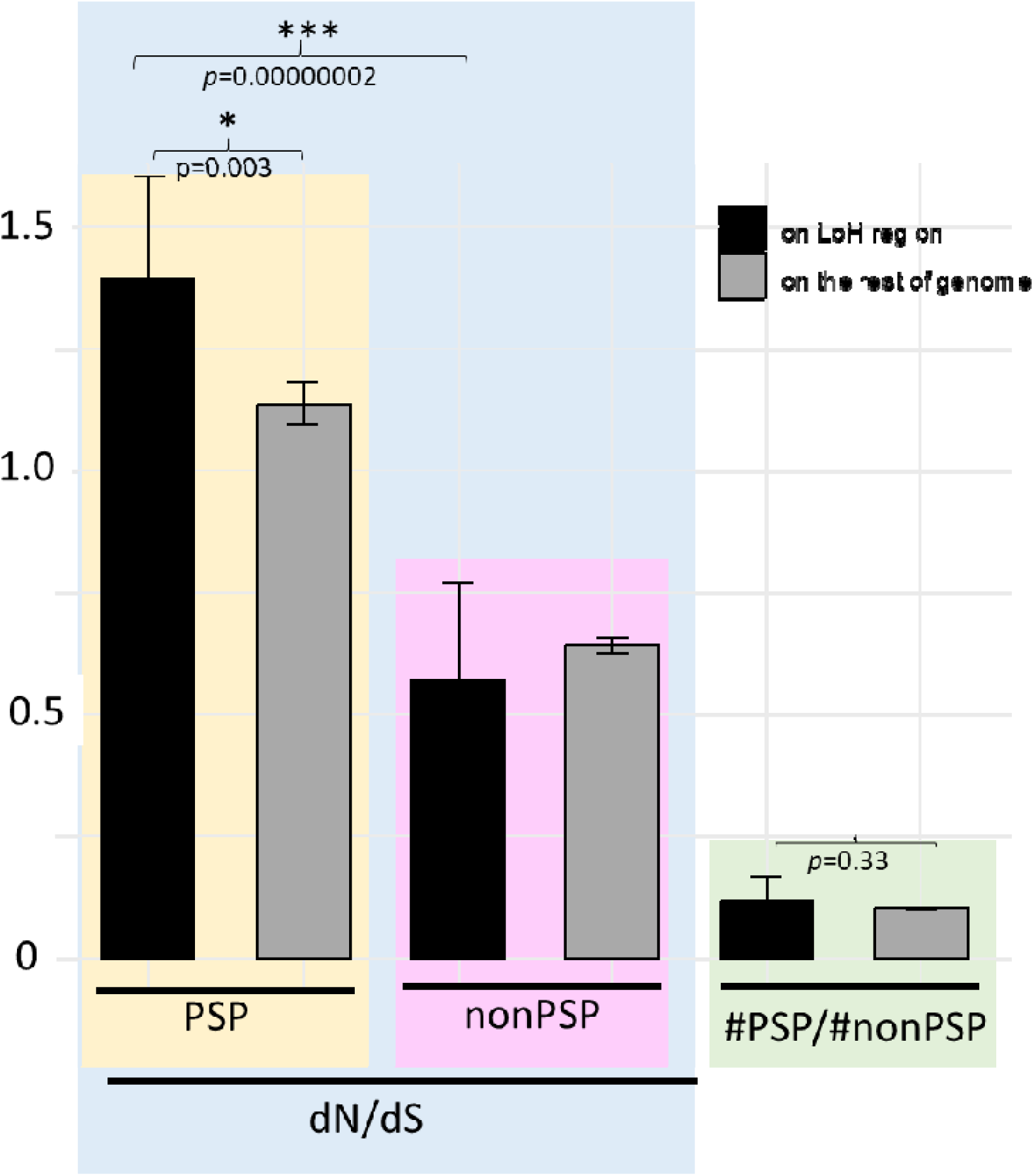
Ratio of non-synonymous to synonymous substitutions (dN/dS) for putative secreted proteins and other protein-coding genes located in LoH regions and in comparison with those in the rest of the genome for each isolate. Abbreviations: *PSP*: putative secreted protein-coding genes, non*PSP*: non-secreted protein-coding genes, #PSP/#nonPSP: ratio of *PSP* to non *PSP*, dN/dS: ratio of non-synonymous to synonymous substitutions. Yellow background color: ratio of non-synonymous to synonymous substitutions in putative secreted proteins located on LoH regions and the rest of the genome. Pink background color: ratio of non-synonymous to synonymous substitutions in other non-secreted protein-coding genes located on LoH regions and the rest of the genome. Green background color: ratio of the number of putative secreted proteins to non-secreted protein-coding genes in LoH and in the rest of the genome. *Significant difference at level *P* < 0.05, ***significant difference at level *P* < 0.001.

Visual inspection of read mapping on the two distinct haplotypes of 18 pairs of curated contigs (354 kb; see Supplementary Method 2) identified 20 LoH events across ten pairs of contigs among nine isolates (for more details, see Supplementary Results 2). These findings confirm the reliability of LoH representation in the *Mg* genomes.

### 2.3 Evidence for meiotic recombination events in *M. graminicola*

On 693,013 SNVs among the 13 *Mg* isolates, 44,527 were retained after eliminating the loci which showed the heterozygosity variants “0/1” in all isolates. Then, in order to test the occurrence of meiotic homologous recombination, we performed a LD analysis as well as a 4- gamete test on the genome-wide SNVs matrix. LD analysis revealed a logarithmic-exponential downward trend in LD between markers as a function of inter-marker physical distance (Figure 4). The LD curve (r^2^) started at high value (0.98), followed by a dramatic slope down close to 0 (r^2^ = 0.125) at 20 kb and which remains at this threshold beyond (Figure 4). In the 4-gamete test, we observed a rapid and exponential increase in the proportion of markers passing the 4-gamete test to reach a value of 0.99 as soon as markers are more than 20 kb away. By performing this same test but using only the 40 fixed single nucleotide mutations observed among the whole genome of isolates (See Supplementary Method 1) the same LD and 4-gamete profiles were observed (Figure S4). The drop of LD and the concomitant increasing proportion of bi-allelic markers passing the 4-gamete test with inter-marker distance constitute strong signatures for meiotic recombination.

**Figure 4.**
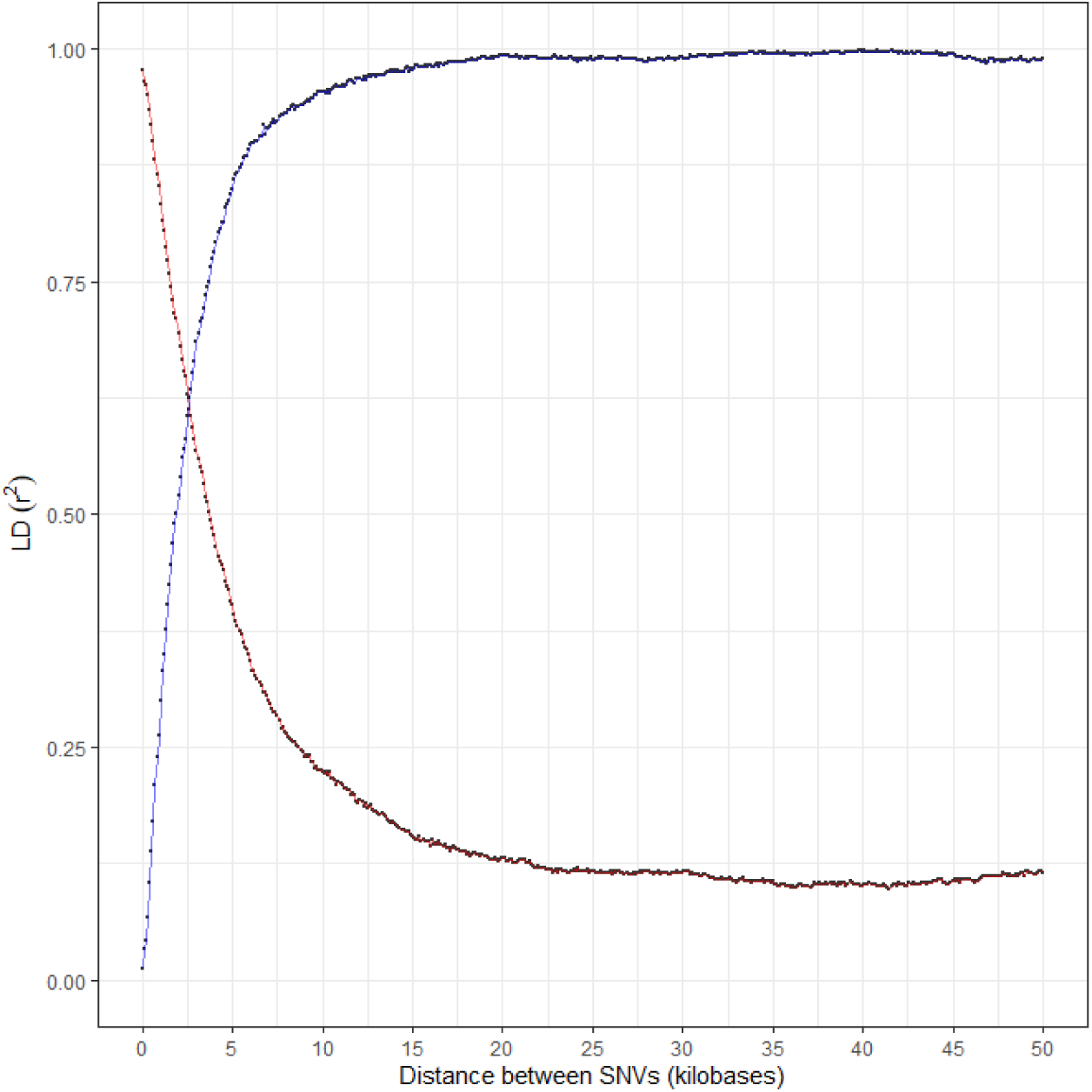
Linkage disequilibrium (LD) between single nucleotide variants (SNVs) as a function of their distance along the genome and 4-gamete test for the 13 *Mg* isolates using the 44,527 SNVs matrix. *The red line indicates r^2^ correlation between markers, indicating LD for each physical distance between DNA polymorphisms. The blue line indicates the proportion of pairs of bi-allelic markers that pass the 4*-*gamete test for each physical distance between the markers on a given scaffold*.

### 2.4 Limited accumulation of single nucleotide variants and no clear geographical pattern of diversity

Clustering of the 13 isolates according to a PCA on the 44,527 SNVs, yielded groups similar to those revealed in the dendrogram based on LoH profiles. A first cluster included the four isolates of Group 1 (Mg-VN18, Mg-VN11, Mg-L2, and Mg-Brazil), while a second group comprised the seven isolates of Sub-group 2.2 (Mg-Bali, Mg-Borneo, Mg-C21, Mg-Java2, Mg-VN6, Mg-VN27, and Mg-L1; Figure 5). In contrast, Mg-C25 and Mg-P (previously defined as Sub-group 2.1 according to LoH profiles) appeared quite distinct from each other and all other isolates. However, the fixation index (*F*_ST_) between Sub-groups 2.1 and 2.2 was very low (mean value: 6.9 × 10□□, weighted value: 9.0 × 10□³), indicating strong genetic connections (Table 3). The *F*_ST_ values between Group 1 and Sub-groups 2.1 or 2.2 were higher, with mean values of 4.2 × 10□² and 1.3 × 10□², respectively. The *F*_ST_ values showed a clear distinction between Group 1 and Group 2, however, those *F*_ST_ values were all below 0.05, indicative of a low genetic differentiation between these groups of isolates (Table 3). Finally, in addition to the analyses of LoH profiles, SNVs also revealed no genetic structuring related to the geographic distribution of *Mg* isolates.

**Figure 5.**
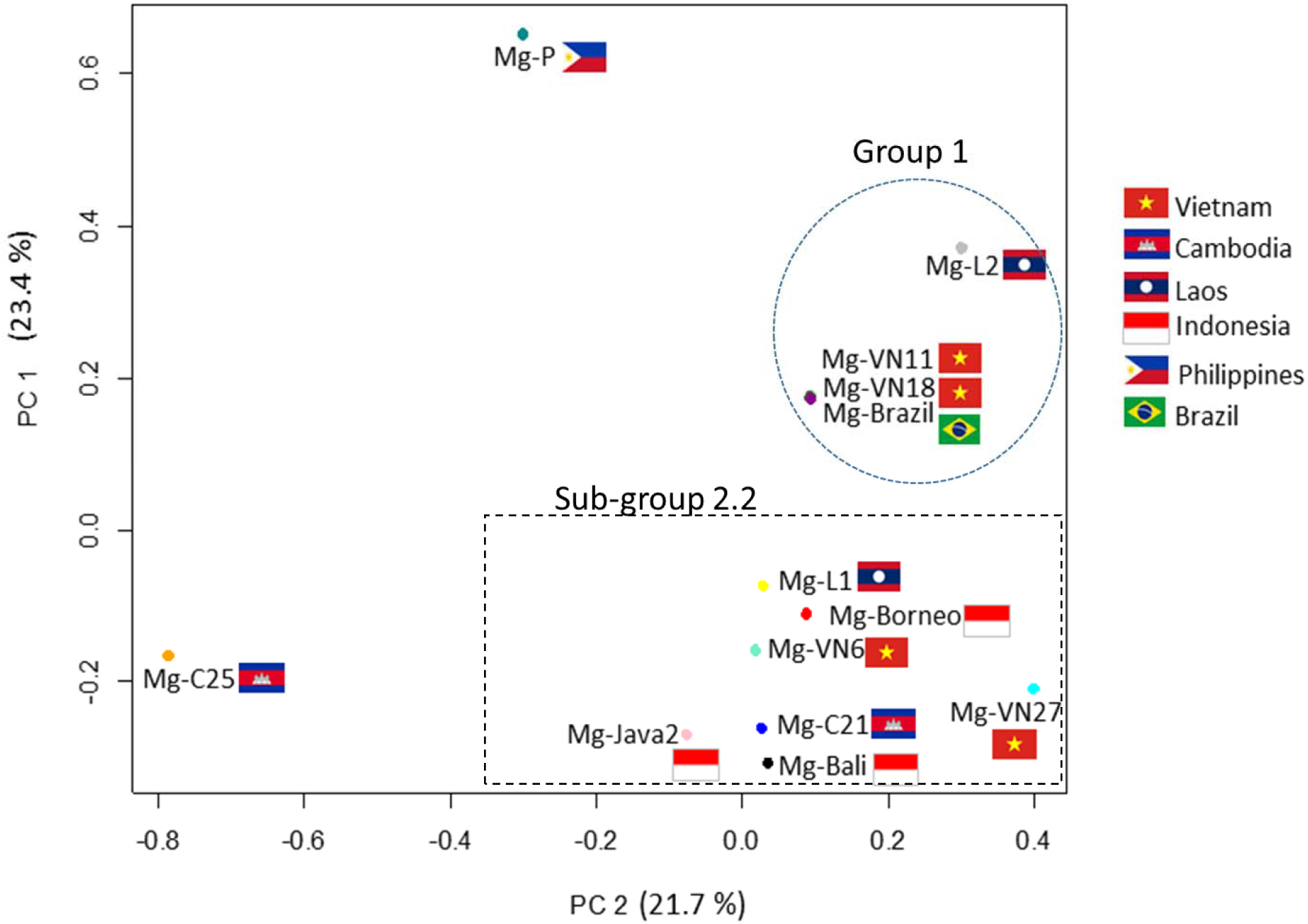
Principal component analysis of the 13 *M. graminicola* isolates using 44,527 SNVs.

**Table 3.** Fixation index (*F*_ST_) based on 44,527 SNVs among groups of isolates. (mean values in bold, weighted values in italic).

|  | Group 1 | Sub-group 2.1 | Sub-group 2.2 |
| --- | --- | --- | --- |
| Group 1 |  | **0.042262** | **0.013906** |
| Sub-group 2.1 | *0.047463* |  | **0.00068335** |
| Sub-group 2.2 | *0.012669* | *0.0089952* |  |

Synonymous and nonsynonymous SNVs were identified in 2,115 protein-coding genes, including 219 genes predicted to encode secreted proteins (Table 4). The nucleotide diversity at synonymous (π_s_) and nonsynonymous (π_n_) sites, along with the π_n_/π_s_ ratio as an indicator of selection efficiency, was calculated for the 13 isolates across protein-coding genes and secreted protein-coding genes at the whole-genome level (Table 4A). The π_n_, π_s_, and π_n_/π_s_ values were generally consistent between the whole gene set and *PSP* genes, although the π_n_/π_s_ ratio was slightly higher in the *PSP* set (0.293 compared to 0.203; Table 4). In comparison with other species, the π_n_ of *M. graminicola* was slightly higher than that of *M. incognita* but marginally lower than the values of the two *Caenorhabditis* species. However, the π_s_ value of *M. graminicola* was ca. twice higher than for *M. incognita* but an order of magnitude lower than those of *C. doughertyi* and *C. brenneri*. Consequently, the π_n_/π_s_ ratio of *M. graminicola* was higher than in *M. incognita* and approximately five times higher than the ratios observed in the two *Caenorhabditis* species (Table 4A).

**Table 4.**
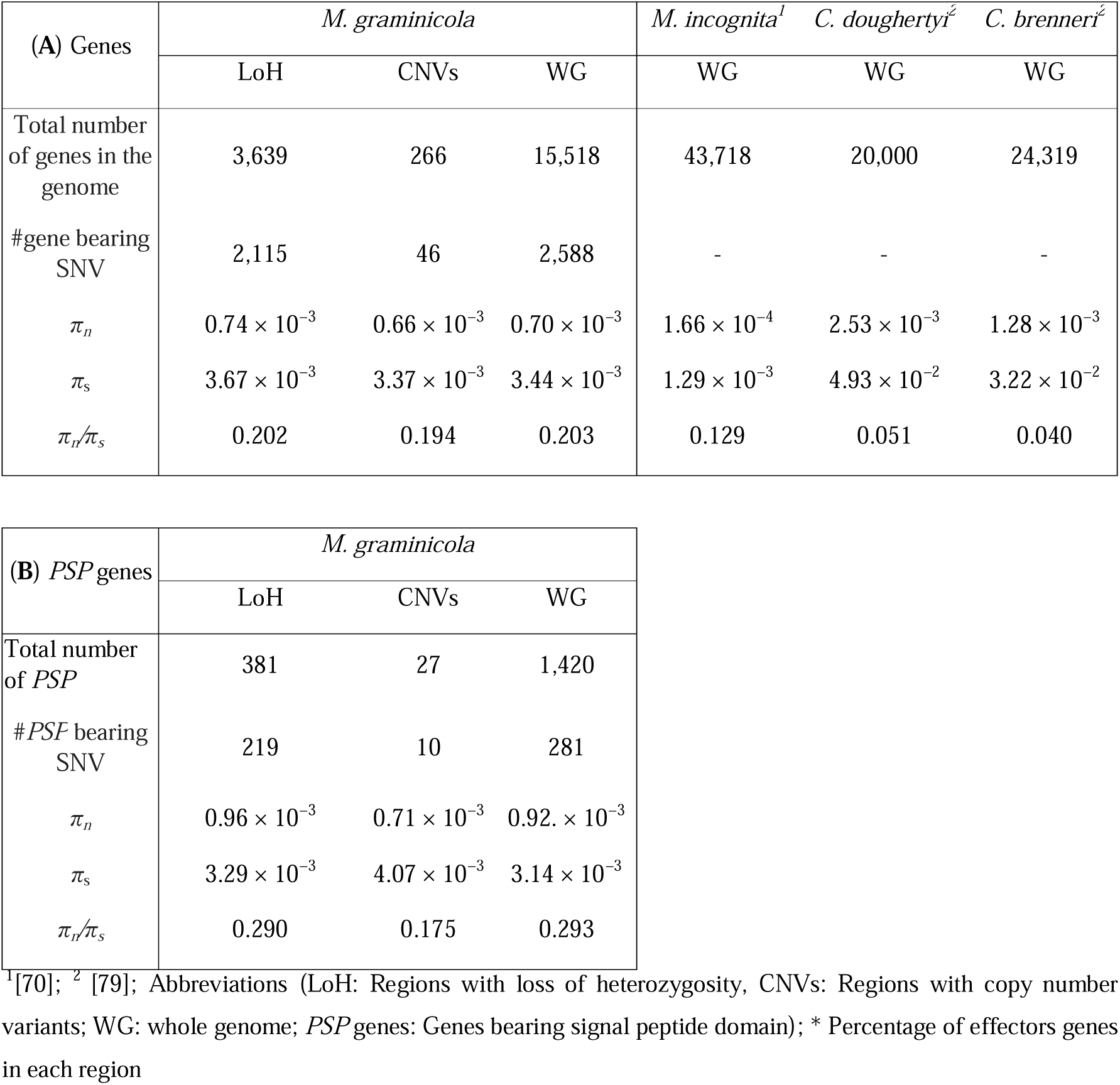
Nucleotide diversity at synonymous (π_s_) and nonsynonymous (π_n_) sites and the π_n_/π_s_ ratio in the coding sequences of all genes (A) and genes bearing signal peptide domains (B) in specific regions of the *M. graminicola* genomes.

| (A) Genes | *M. graminicola* |  |  | *M. incognita*<sup>1</sup> | *C. doughertyi*<sup>2</sup> | *C. brenneri*<sup>2</sup> |
| --- | --- | --- | --- | --- | --- | --- |
|  | LoH | CNVs | WG | WG | WG | WG |
| Total number of genes in the genome | 3,639 | 266 | 15,518 | 43,718 | 20,000 | 24,319 |
| #gene bearing SNV | 2,115 | 46 | 2,588 | - | - | - |
| $\pi_n$ | $0.74 \times 10^{-3}$ | $0.66 \times 10^{-3}$ | $0.70 \times 10^{-3}$ | $1.66 \times 10^{-4}$ | $2.53 \times 10^{-3}$ | $1.28 \times 10^{-3}$ |
| $\pi_s$ | $3.67 \times 10^{-3}$ | $3.37 \times 10^{-3}$ | $3.44 \times 10^{-3}$ | $1.29 \times 10^{-3}$ | $4.93 \times 10^{-2}$ | $3.22 \times 10^{-2}$ |
| $\pi_n/\pi_s$ | 0.202 | 0.194 | 0.203 | 0.129 | 0.051 | 0.040 |

| (B) PSP genes | *M. graminicola* |  |  |
| --- | --- | --- | --- |
|  | LoH | CNVs | WG |
| Total number of PSP | 381 | 27 | 1,420 |
| #PSP bearing SNV | 219 | 10 | 281 |
| $\pi_n$ | $0.96 \times 10^{-3}$ | $0.71 \times 10^{-3}$ | $0.92 \times 10^{-3}$ |
| $\pi_s$ | $3.29 \times 10^{-3}$ | $4.07 \times 10^{-3}$ | $3.14 \times 10^{-3}$ |
| $\pi_n/\pi_s$ | 0.290 | 0.175 | 0.293 |

<sup>1</sup>[70]; <sup>2</sup> [79]; Abbreviations (LoH: Regions with loss of heterozygosity, CNVs: Regions with copy number variants; WG: whole genome; PSP genes: Genes bearing signal peptide domain); \* Percentage of effectors genes in each region

### 2.5 Independent accumulation of copy number variants among isolates

Comparative genome analysis of the 13 isolates revealed 92 CNV regions across 39 scaffolds, with CNV sizes ranging from 3 to 30 kb (Tables S3 and S5). While Mg-Bali, Mg-Java2, Mg- VN27, Mg-P, Mg-C21, and Mg-C25 exhibited more deletions than duplications, other isolates (Mg-VN18, Mg-VN6, Mg-VN11, Mg-Borneo, Mg-Brazil, Mg-L1, and Mg-L2) tended to accumulate more duplicated copies (Figure 6A).

**Figure 6.**
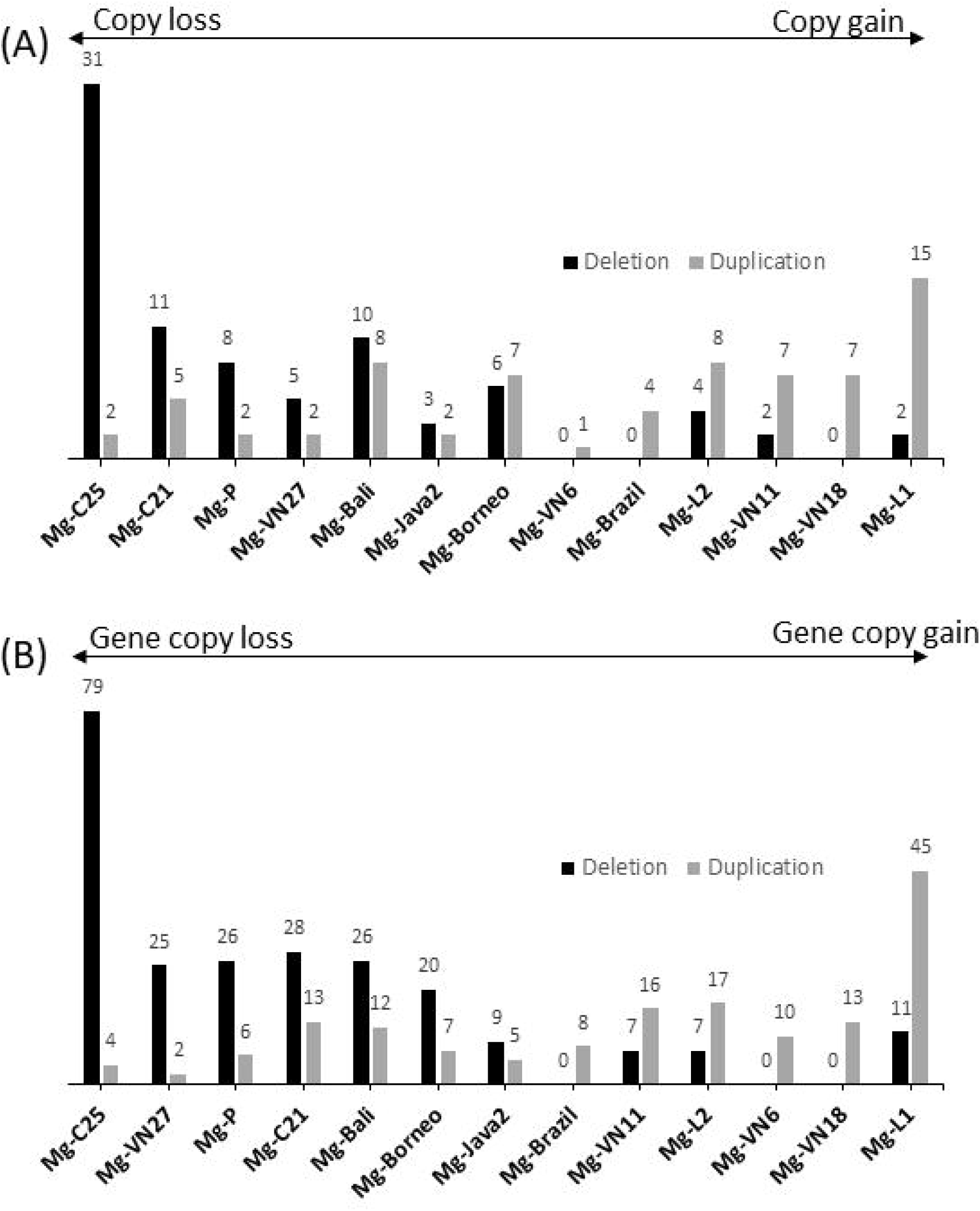
Copy number variants (CNVs) among the 13 *M. graminicola* isolates: (A) Number CNVs regions experiencing gains or losses their copies among isolates, and (B) Number of genes affected by CNVs regions resulting gain or loss of gene copies among isolates.

Interestingly, four isolates (Mg-VN18, Mg-VN11, Mg-Brazil, and Mg-L2) that have accumulated more duplications also had fewer LoH events and fewer point mutations (Figure 6A, Table 2). Conversely, isolates that accumulated more deletions (Mg-P, Mg-C21, and Mg- C25) showed a higher number of LoH events (Figure 6A, Table 2) and a greater number of point mutations among the fixed 40 SNVs (Supplementary Results 1). Additionally, detailed analysis of read coverage across the 18 pairs of homologous contigs indicated that the ACC4 haplotype was lost in Mg-C25 (Figure S7). We observed overlapping of LoH events, duplications and deletions on several scaffolds (e.g. mg19, mg71, mg75, mg76, mg77, mg78, mg85; Figure S9).

Overall, alignment of the CNV regions (652.4 kb) with the reference genome [33] indicated that 266 genes might be affected by CNVs. The Mg-C25 isolate is notably distinguishable from the others due to a significant number of gene copy losses (79 genes; Figure 6B). Five isolates, Mg-L2, Mg-C21, Mg-P, Mg-VN27, and Mg-Java2, exhibited more deleted genes than duplicated genes (Figure 6B). In contrast, isolates Mg-L1, Mg-VN18, Mg- VN6, Mg-VN11, and Mg-Brazil tended to have more duplicated genes (Figure 6B). A total of 27 *PSP* genes were affected by CNVs. Among these, ten genes experienced complete deletion (loss of both copies), 11 genes underwent partial deletion (loss of one copy), and six genes were duplicated, resulting in three or four additional copies (Table S3). Interestingly, some *PSP* genes encoding enzymes such as pectate lyase, peptidase and glycosyltransferase, which are likely involved in plant cell wall and protein degradation, were deleted or reduced in copy number (Table S4). Meanwhile, *PSP* genes potentially involved in inhibiting host plant proteases (e.g., Zona pellucida domain, Serine protease inhibitor-like) and feeding site formation (e.g., LRP chaperone MESD, EGF-like domain) were duplicated (Table S4). In CNV regions, synonymous and non-synonymous SNPs were identified in 46 protein-coding genes, including 10 *PSP* genes. Interestingly, isolates Mg-C25 and Mg-VN27 had deleted copies of all 10 *PSP* genes. The ratio of non-synonymous to synonymous substitutions (dN/dS) was less than 1 for both protein-coding genes (dN/dS = 145/151 = 0.96) and *PSP* genes (dN/dS = 43/55 = 0.78; data not shown). Besides, the selection efficiency ratio (π_n_/π_s_) was 0.197 for protein-coding genes and 0.175 for secreted protein-coding genes within the CNV regions, both of which were lower than the corresponding ratios at the genome level (Table 4).

Visual inspection of read mapping on both haplotypes of 18 pairs of contigs (354 kb; 0.43% of the genome) revealed two cases of haplotype deletion (corresponding to CN1 in Mg-C25 and Mg-Bali) and one case of haplotype duplication (corresponding to CN3 in Mg-Borneo) (for more details, see Supplementary Results 2). These results confirmed the reliability of CNVs representation in the genome of *Mg*.

## 3. DISCUSSION

### 3.1 Homologous recombination suggests occasional sexual reproduction but prevalence of meiotic parthenogenesis

This study allows us to assess the extent of genomic variation in the nuclear genome of *M. graminicola* during its recent diversification. At the same time, our analysis of SNVs revealed a rapid increase in the proportion of marker pairs passing the 4-gamete test, along with a decrease in linkage disequilibrium (LD) as the physical distance between markers increased. Such a rapid decline in LD in *M. graminicola* should be due to homologous recombination as it has been reported in the Bdelloid rotifer *Adineta vaga* [26]. This pattern contrasts with the lack of recombination observed in the apomictic *M. incognita* but resembles the findings in the meiotic amphimictic nematode *Globodera rostochiensis* [70]. While translocations and gene conversion can break down LD [80], they would not produce the logarithmic-exponential relationship we observed between inter-marker distance and the proportion of bi-allelic markers passing the 4-gamete test. Instead, short regions of loss of heterozygosity (LoH), often referred to as interstitial LoH, were randomly distributed across the scaffolds. In contrast, large LoH regions (>100 kb) were frequently found at scaffold ends, indicating terminal LoH, likely resulting from reciprocal crossover events or break-induced repair during cell division [81, 82].

The heterozygosity level in the *M. graminicola* genome ranged from 1.78 to 3.43% among 13 isolates (at *k*-mer = 21 bp), consistent with other meiotic and sexually reproducing animals [83]. This heterozygosity is higher than that observed in species with an intraspecific origin of parthenogenesis (e.g., *Anthocomus rufus*, *Ooceraea biroi*; 0.03–0.83%) but falls within the range of some hybrid-origin parthenogenetic species (e.g., *Poecilotheria formosa*, *Panagrolaimus davidi*, *Diploscapter coronatus*; 1.43–4%) [27]. However, it remains much lower than the 7-8% observed between subgenomes in polyploid hybrids within the *Meloidogyne* genus [35, 77].

Three hypotheses may explain the observed heterozygosity level in *M. graminicola*. First, this species might have originated from a previous homoploid hybridization with higher heterozygosity, which has since decreased due to genetic erosion mechanisms like recombination and gene conversion. Phylogenetic analysis of low-copy nuclear regions suggests a potential hybrid origin related to *M. oryzae* [15]. Second, if *M. graminicola* undergoes meiotic parthenogenesis via central fusion, the separation of homologous chromosomes during meiosis would preserve initial heterozygosity, which is advantageous for adapting to environmental challenges [84]. In contrast, terminal fusion could lead to rapid genetic diversity loss, as observed in *M. hapla* [85]. Third, the observed heterozygosity might arise from sexual reproduction between closely related individuals [86]. However, the shared LoH profile (∼1.1% of genome size) among nine isolates suggests primarily clonal reproduction, with occasional sexual events allowing for the conservation of specific heterozygosity patterns across isolates.

The occurrence of meiotic homologous recombination, indicative of sexual reproduction, is strongly supported by the results of the 4-gamete test, LD pattern analysis, and observed LoH in *M. graminicola*. Interestingly, while the increase in markers passing the 4-gamete test and decrease in LD occur over relatively short distances (<1 kb) in *G. rostochiensis* [70], in *M. graminicola*, this decline occurs over a greater distance (∼20 kb). This suggests that recombination, although present in *M. graminicola*, occurs less frequently compared to *G. rostochiensis*, likely due to its predominant mode of meiotic parthenogenesis. Triantaphyllou’s [31] cytogenetic analyses support this conclusion, indicating that *M. graminicola* primarily reproduces via meiotic parthenogenesis.

### 3.2 Recombination and loss of heterozygosity contribute to genomic variation in *M. graminicola* and are associated with positive selection

Meiotic recombination and LoH could provide genomic plasticity for adaptive evolution in *M. graminicola*. Meiotic recombination might create a chance to obtain new gene combinations that are favorable for parasitism. Even in primarily clonal or parthenogenetic species, occasional recombination through rare sexual events can introduce genetic diversity, enabling populations to adapt to changing environments and resist selective pressures [87, 88]. For instance, in the facultative parthenogenetic root-knot nematode *Meloidogyne hapla*, meiotic recombination has been shown to promote genetic diversity, which helps the species to adapt to diverse hosts and environmental conditions [89]. Similarly, in the asexual fungal parasite *Candida albicans*, recombination events, even in the absence of full sexual cycles, introduce genetic variations that contribute to its adaptability, particularly in response to antifungal drugs [90].

Loss of heterozygosity is expected in most forms of meiotic parthenogenesis [91, 92]. It might unmask recessive alleles that may confer advantageous traits. In clonal or asexual populations, LoH events can lead to genomic regions becoming homozygous, which can accelerate adaptation by allowing beneficial alleles to be fully exposed to selection. This mechanism has been observed in other parthenogenetic species, where LoH contributes to adaptability in varying environments. For example, in *Daphnia pulex*, LoH events enable adaptation to environmental stresses by revealing hidden beneficial alleles, thus increasing the fitness of populations [93]. Similarly, in the plant-parasitic fungus *Candida albicans*, LoH facilitates adaptation by promoting drug resistance, as specific LoH events lead to the expression of resistance-conferring mutations [94]. Bdelloid rotifers seem to survive extreme conditions (e.g. freezing, deep vacuum, UV and high doses of ionizing radiation) despite its asexual reproduction (“evolutionary scandal”) thanks to effective antioxidants and double-strand breaks (DSBs) repairing mechanisms, which are often repaired through homologous recombination processes that can further promote LoH [95–97]. LoH potentially unmasks recessive alleles that enhance virulence or allow adaptation to different host environments [98]. In *M. graminicola*, LoH regions covered up to 7% of the genome, with variation among isolates, and impacted a number of putative secreted protein-coding genes (*PSP* genes). The ratio of non-synonymous to synonymous substitutions (dN/dS) for *PSP* genes in LoH regions was greater than 1 (1.40 ± 0.20), indicating positive selection [99]. Notably, this dN/dS ratio was higher than that observed for *PSP* genes in the rest of the genome, suggesting that *PSP* genes within LoH regions are under stronger positive selection or adaptive pressure compared to those located elsewhere. Positive selection on *PSP* genes could reflect an adaptive response to host defenses, enabling the nematode to better evade immune responses or exploit the host. Overall, genome recombination/reorganizations including LoH can drive evolutionary change, potentially enhancing the resilience and adaptability of parthenogenetic parasites like *M. graminicola* in response to environmental pressures.

### 3.3 Copy number variation as a mechanism allowing potential adaptation

CNVs have been identified in all areas of life, from bacteria, to plants and animals and appear to play important functions in the evolution of genomes [29, 100]. CNVs consist of DNA segments that are variable in copy number and dispersed throughout the genome [101]. CNVs can either be inherited from the previous generation or appear *de novo* through duplication/deletion events, and their fixation by drift or selection may contribute to the creation of genetic novelty resulting in species adaptation to stressful or novel environments [102, 103].

In this study, we detected numerous CNVs (∼1.6% genome length) in all *M. graminicola* isolates. Most of these CNVs appeared quite recently and independently among isolates without geographical relationship. It is likely that each isolate itself accumulates gene copy gain or loss while adapting to the environment and the host. The overall π_n_/π_s_ (less than 0.2) and dN/dS (less than 1) ratios in CNV regions suggest purifying selection, which may act to conserve essential gene functions while allowing occasional innovation through CNV-mediated gene duplication or loss [104]. We observed that certain isolates (eg. Mg-C25, Mg-VN27, Mg-P, Mg-C21) exhibited significant gene loss events and tended to have a higher proportion of homozygous nuclear regions. Gene loss may be linked to chromosomal rearrangements, such as deletions or duplications, which can also affect heterozygosity. This explains also for overlapping regions on genomic scaffolds between LoH regions and CNVs (Figure S9). LoH might also result in a deletion of one gene copy recognized by the plant’s immune system, allowing resistance to be overcome [30]. The prevalence of deletions in *PSP* genes, such as pectate lyase and peptidase, may indicate a trend toward genome streamlining in *M. graminicola*, where non-essential or redundant genes are eliminated to reduce metabolic costs. Additionally, losing one gene copy may provide an advantage by helping the organism evade recognition by the plant’s immune system [29]. This could be advantageous in environments where these functions are unnecessary or where alternative pathways compensate for the loss. The duplication of *PSP* genes related to host defense inhibition and feeding site formation likely reflects positive selection for traits that enhance virulence and adaptability to host defenses. This aligns with the arms race model of host-pathogen coevolution [105]. The distinct CNV patterns in Mg-C25 and Mg-VN27 suggest that CNVs contribute to isolate-specific adaptation.

Recent studies also detected CNVs in Cyst and RKN plant parasitic nematodes [30, 106, 107]. Some of these CNVs seem to be correlated with the ability of the parasite to evade the defense mechanisms and to become more virulent [30, 106, 107]. Interestingly, convergent gene loss events have been associated with virulent isolates of *M. incognita* [30]. This correlation does not yet point to causation, but it does raise questions about the potential role of CNVs in the acquisition of virulence. The genetic variations generated by these CNVs could reflect an adaptive response to different environments and in particular to the host plant, in which this obligate parasite must establish a compatible interaction to be able to reproduce. A virulent population of *M. graminicola* was recently discovered in South-East Asia [108]. For example, it would be interesting to find out whether CNVs are specifically associated with this virulent population, as already observed in *M. incognita* [30].

### 3.4 Recent spread of *M. graminicola* and potential impact of clonal reproduction

In organisms with a short generation time and relatively small genomes, recombination defects could lead to the isolation of a large set of mutants [109]. Surprisingly, in *M. graminicola*, single nucleotide mutations appear to be fixed at a low frequency. The search for such mutations is challenging on the whole genome, because of its heterozygous status, the presence of LoH, and the fact that only the haploid reference is available. The 40 selected SNVs (in total of 44,527 SNVs) do not represent the total diversity of isolates, but only a subset of SNVs that were not linked to the genome rearrangement events among isolates in this study (see Supplementary Result 1). While no sequence polymorphism was previously reported in nuclear regions covering 40.6 kb [15], we detected only two SNVs in 354 kb, allowing us to estimate that the average SNV spanned 2/354 kb = 0.0005% of the nucleotides (see Supplementary Result 1). Even though mitochondrial diversity is low, the SNV percentage (0.07%) is one hundred times higher than the one of the nuclear genome in *M. graminicola* [15], which is consistent with that observed in many species of the phylum Nematoda and especially in *Meloidogyne* species [70, 110, 111]). In *M. incognita* populations, SNVs account for 0.02% of the nucleotides in the nuclear genome, which is even higher than that in *M. graminicola* [70]. The low level of polymorphisms between isolates indicate SNVs might not be driving force of genome evolution in *M. graminicola*. It would be especially interesting to explore the molecular machinery of *M. graminicola*, as there appears to be a particularly efficient DNA repair process. Alternatively, it is possible that *M. graminicola* underwent a recent population expansion from a common ancestral isolate.

PCAs based on SNVs and LoH patterns revealed two main groups with a lack of geographical structuring of isolates as already reported for mitochondrial genomes and for *M. incognita* [15, 16, 28, 70]. Isolates in Group 1 (Mg-VN18, Mg-VN11, Mg-L2, and Mg-Brazil) were genetically similar and showed more genetic differentiation (higher *F*_ST_ value) from Group 2. However, the *F*_ST_ values remain below 0.05, signifying low overall genetic divergence. Sub-group 2.2 forms a larger cluster with seven isolates that are closely related and limited differentiation with extremely low *F*_ST_ values between isolates (mean: 6.9 × 10□□). This sub-group may represent a more recently diversified lineage that expanded across Southeast Asia, as suggested by the presence of isolates from Indonesia (Bali, Borneo, Java) and Vietnam. Although previously placed within Group 2 based on LoH profiles, Mg- C25 and Mg-P show unique genetic characteristics in SNV analyses, appearing distinct from both Sub-group 2.2 and Group 1. These isolates may represent transitional or hybrid lineages, or they could be the result of local adaptation or reduced gene flow from the other groups. Alternatively, they might originate from a completely different genotype or geographical region than the rest of the isolates. Group 1 may represent an older, genetically distinct lineage that diverged early from Group 2, while this latter could be a younger lineage or an actively exchanging population.

In addition, other molecular diversity among isolates, such as LoH and CNVs, also showed no clear association with the geographic origin of isolates. Furthermore, the small genetic distance between Southeast Asia populations suggests they recently derived from each other. These observations reinforce the hypothesis of a recent expansion of *M. graminicola* isolates over Southeast Asia and probably globally. This expansion could have been driven by agricultural practices, such as the spread of rice varieties across the region. It is likely that these isolates were recently spread from a common founder population and then accumulated molecular diversity (SNV, LoH, and CNVs). During the adaptation process, some isolates (Mg-C25, Mg-C21, Mg-P, and Mg-VN27) have accumulated more mutations than others. Interestingly, haplotype networks using mutations in mitogenome (11 SNVs, 5 InDels, 1 inversion) and nuclear genome (40 SNPs) among 13 isolates showed strong correlation, suggesting a common direction of molecular evolution of both mitogenome and nuclear genome accompanying the adaptation to new environments. These mutations may have accumulated independently following the expansion across a large area.

The elevated π_n_/π_s_ ratio observed in *M. graminicola* compared to those in *M. incognita*, *C. doughertyi* and *C. brenneri* suggests a relatively higher level of nucleotide diversity at nonsynonymous sites compared to synonymous sites [112]. This could indicate that *M. graminicola* is experiencing adaptive evolution, where nonsynonymous mutations are subject to positive selection or are less constrained functionally [113]. This apparent paradox, where meiotic parthenogenesis in *M. graminicola* exhibits lower efficacy of selection than mitotic parthenogenesis in *M. incognita*, may stem from the fundamental differences in the genetic mechanisms underlying the two reproductive modes. Meiotic parthenogenesis involves recombination during gamete formation, which generally promotes the purging of deleterious mutations, preventing Muller’s ratchet [114]. In contrast, mitotic parthenogenesis, by lacking recombination, maintains permanent linkage among mutations, thereby impeding the selective purging of deleterious alleles and facilitating their gradual accumulation over time [27]. Furthermore, facultative parthenogenesis, as observed in *M. graminicola*, may introduce additional complexities due to occasional sexual reproduction, which can disrupt linkage disequilibrium and dilute the effects of selection [115, 116]. A study by Romiguier et al. [79] examining 17 species—ten obligate sexual species (e.g., seahorse, tortoise, sea urchin) and seven facultative parthenogenetic species (e.g., ants, termites)—revealed that π_n_/π_s_ ratios ranged from 0.21 to 0.37. These values, which were higher than that in *M. graminicola*, suggest an even lower efficacy of selection in these sexual/parthenogenetic species than in the facultative parthenogenetic species *M. graminicola*. Therefore, the efficacy of selection may not always be determined solely by the mode of reproduction, but rather by the specific genetic mechanisms (such as recombination rates, mutation accumulation, genetic drift, and the presence of other evolutionary pressures) at play within the genome.

## 4. CONCLUSION

This study presents an improved genome annotation for the *Meloidogyne graminicola* reference genome, providing a crucial foundation for advanced genetic analyses. A key highlight is the first comprehensive insight into the evolutionary traits of this meiotic root-knot nematode through population genomics. For the first time, meiotic recombination has been confirmed within *M. graminicola* populations, shedding light on its reproductive dynamics. Key evolutionary mechanisms, including loss of heterozygosity (LoH) and copy number variation (CNV), were explored among isolates, revealing their potential impact on secreted protein-coding genes and underscoring their role in adaptive evolution. The analysis of single nucleotide variants (SNVs) and LoH patterns identified two distinct clusters of isolates, likely diverging from a common ancestor. Moreover, the low genetic differentiation among populations supports a scenario of recent radiation and rapid spread across Southeast Asia. Our data also indicate that *M. graminicola* primarily reproduces clonally, with occasional sexual reproduction, combining the advantages of both modes to enhance its adaptability and evolutionary potential as a parasite. Although further experiments and analyses are necessary to document the relative importance of each reproductive strategy, our study provides valuable genomic resources and new insights. These findings will be instrumental for future in-depth investigations aimed at understanding the evolutionary dynamics and adaptive mechanisms driving the success of this parasitic nematode.

## 5. MATERIALS & METHODS

### 5.1. Nematode sampling, DNA extraction, sequencing and reads cleaning

Thirteen isolates of *M. graminicola* originating from distant locations were collected in rice fields in South East Asia and Brazil (Table S1; [15]). These isolates were obtained after single nematode infection [12]. Subsequently, each isolate was propagated for two months on the susceptible *Oryza sativa indica* var. IR64 before DNA extraction [15]. Briefly, juveniles and eggs were extracted from roots using a hypochlorite extraction method and a blender, then treated for 15 min in 0.8% hypochlorite at room temperature, purified on a discontinuous sucrose gradient, and finally rinsed several times with sterile distilled water before being stored at −80°C. The total genomic DNA (gDNA) was isolated after treatment with proteinase K and extraction with phenol-chloroform followed by ethanol precipitation [36]. DNA quality and quantification were assessed with a NanoDrop spectrophotometer (Thermofisher, Waltham, MA, USA) and PicoGreen^®^dsDNA quantitation assay (Thermofisher). Finally, DNA integrity was checked by electrophoresis, loading 1 µL on a 1% agarose gel.

High□depth short□read sequencing was performed at the GeT□PlaGe core facility, INRAE Toulouse. For each accession, 300 ng of double-stranded DNA (dsDNA) was used for shotgun sequencing using Illumina technology (San Diego, CA, USA). DNAs were sonicated to get inserts of approximately 380 bp for adaptor ligation using the Illumina TruSeq DNA Sample Prep v2 kit. Libraries were multiplexed and inserts were then sequenced from both ends (2×100 bp, 2×125 bp, 2×150 bp) on HiSeq 2000 or 2500 (Illumina) (Table S1).

Illumina sequences were trimmed and cleaned. First, adapter sequences were removed with cutadapt [37]. Then, overrepresented sequences and low quality bases (Q < 20) were removed with Skewer [38]. Subsequently, the quality of sequences was assessed with fastQC [39]. Minor GC-rich peaks (ca. 70%) were revealed in five isolates (Mg-Bl, Mg-Bn, Mg-C21, Mg-C25, and Mg-P) indicating potential contaminants (e.g. from microorganisms). Sequences of such contaminants were detected and removed based on the method proposed by Kumar et al. [40]. In short, the genome of each isolate was pre-assembled in contigs using Spades [41], that were blasted against the NCBI nt database to annotate taxa with high identity (>75%, e-value = 1e-25). Finally, the sequencing coverage, GC content, and length, as well as the annotated taxa were summarized and visualized on each pre-assembled contig using blobtools [42]. Based on this information, contaminations of Proteobacteria and Firmicutes were detected only in these five isolates. Therefore, reads from these contaminants were removed using Bowtie2 [43], resulting in the final set of cleaned reads. Five isolates (Mg-Bali, Mg- Borneo, Mg-Java2, Mg-L2, Mg-VN18) showed high genomic coverage (288 to 568× relative to the size of the reference haploid genome - 41.5 Mb), while other isolates had lower coverage ranging from 21 to 39×. Therefore, genomic sequences of those five isolates were down-sampled to 20% using the SEQTK tool [44].

### 5.2. Gene annotation and secreted proteins prediction for the reference genome sequence of *M. graminicola*

The whole genome sequence of *M. graminicola* published by Phan et al. [33] was used as reference. Detection of gene models was done with the fully automated pipeline EuGene-EP v1.6.3 [45]. EuGene has been configured to integrate similarities with known proteins of *Caenorhabditis elegans* (PRJNA13758), a previously published version of the *M. graminicola* predicted proteome [46], both downloaded from Wormbase ParaSite [47], as well as the “nematoda” section of UniProtKB/Swiss-Prot library [48], with the prior exclusion of proteins that were similar to those present in RepBase [49].

Three datasets of *M. graminicola* transcribed sequences were aligned to the genome and used by EuGene as transcriptional evidence: (i) de novo assemblies of the *M. graminicola* J2 stage transcriptome generated using Trinity [50], followed by a cleanup step that retains only the transcript with the longest ORF for each Trinity locus; (ii) StringTie [51]; and (iii) Oases [52]. The alignments of datasets on the genome, spanning 30% of the transcript length with at least 97% identity were retained.

The EuGene default configuration was edited to set the “preserve” parameter to 1 for all datasets, the “gmap_intron_filter” parameter to 1, the minimum intron length to 35 bp, and to allow the non-canonical donor splice site “GC”. Finally, the Nematode specific Weight Array Method matrices were used to score the splice sites (available at this URL: http://eugene.toulouse.inra.fr/Downloads/WAM_nematodes_20171017.tar.gz). To assess the completeness of the predicted protein set in this study (15,518 proteins, Eugene_Mg) and the three previous published proteome versions ASM1477313v1 (10,331 proteins; [33]), PRJNA411966 (14,062 proteins; [34]), and T2T (12,968 proteins; [35]), we used BUSCO v5.5.0 [53] in protein mode and “-long” mode for AUGUSTUS optimization. We used three different datasets including eukaryota odb10 (255 groups), metazoa odb10 (954 groups), and nematoda odb10 (3,131 groups).

We also used SignalP5.0 [54] with the option ‘mature’ to predict signal peptides for secretion in the whole set of proteins annotated from the genome and produce fasta sequences with cleaved signal peptides. We then searched for transmembrane helix in these cleaved sequences with Tmhmm2.0c [55]. All proteins with a predicted signal peptide and no predicted transmembrane region were considered possibly secreted.

All predicted proteins were scanned for the presence of conserved protein domains and motifs using InterProScan v5.51-85.0 [56, 57] with the options -iprlookup, -goterms and -pa to assign Gene Ontology (GO) terms and KEGG, MetaCyc and Reactome biochemical pathways based on detection of Interpro domains.

### 5.3. Calling and genotyping nucleotide variants in the whole genome of the 13 *Mg* isolates

To detect variants among the genome sequences of the 13 *Mg* isolates, Illumina reads of each isolate were mapped to the haploid reference genome of *M. graminicola* (isolate Mg-VN18; [33]) using the BWA-MEM software [58]. SAMtools [59] was used to filter alignments with MAPQ lower than 20, to sort the alignment file by reference position, and to remove multi-mapped alignments. GATK [60] was used to mark and remove the duplicated reads (MarkDuplicate). FreeBayes [61] was used to detect the variants (SNVs and short indels) using all alignments simultaneously to produce a variant call file (VCF). The VCF file was filtered using vcftools [62], retaining only the positions that had more than 30 phred-scaled probability (minGQ), a minimum coverage depth (minDP) of ten and no missing data. Given that the *M. graminicola* genome has an average heterozygosity of 1.36% ± 0.78 ([15, 33]) and that the final reference genome is a haploid genomic sequence consisting mainly of collapsed haplotypes [33], we had to establish a specific code for the sequence polymorphisms observed. Thus, after mapping of the Mg-VN18 reads to the Mg-VN18 haploid genomic reference sequence [33], the collapsed contigs obtained corresponded either to heterozygous regions [i.e., two haplotypes (or alleles) detected and coded as follow: genotype “0/1” where 0 = first haplotype sequence identical to the reference, and 1 = second haplotype sequence]; or to homozygous regions (identical to the reference sequence and coded “0/0”). When reads of other isolates were mapped to the reference genome, six possible genotypes were defined as follow: “0/0” = homozygous region with the same sequence as the reference, “0/1” = haplotype of Mg-VN18, “1/1” = homozygous region bearing only the alternative haplotype of Mg-VN18, “0/2” = heterozygous region containing the reference haplotype sequence and a second alternative allele (distinct from this detected in Mg-VN18), “1/2” = heterozygous region bearing first and second alternative haplotypes, and “2/2” = homozygous region bearing only a second alternative allele (not detected in Mg-VN18). A matrix of the variant genotypes of the 13 isolates was constructed allowing us to score polymorphism for each position in the genome. The distribution of variant genotypes for each isolate on the reference genome scaffolds was visualized using CIRCOS (http://circos.ca/) in order to reveal specificities of each isolate (i.e., polymorphism, genome organization). Note that homozygous “0/0” and heterozygous “0/1” genotypes among all isolates represent non-variable regions. Meanwhile, heterozygous “1/2” and “0/2” genotypes as well as homozygous “1/1” and “2/2” genotypes correspond to true variants between the reference Mg-VN18 and any other isolate. Therefore, the final matrix of single nucleotide variants (SNV) only includes loci which have at least one variant genotype (“1/1”, “2/2”, “1/2”, or “0/2”) among all isolates.

We used SnpEff [63] to annotate nucleotide variability among 13 isolates at nonsynonymous and synonymous sites using the default parameters. The function bpppopstats from the Bio libraries [64] was used to estimate the nucleotide variability at nonsynonymous and synonymous sites as well as efficacy of purifying selection (π_n_ /π_s_) in the coding sequence, based on the standard genetic code.

### 5.4. Detection of loss of heterozygosity across the genome of the 13 *Mg* isolates

The heterozygosity of the 13 *Mg* isolates was estimated using HiSeq genome sequences. Jellyfish v2.3.0 [65] was employed to extract and count canonical *k*-mers (*k* = 19, 21, 25, and 27 nucleotides) from cleaned Illumina reads. For each *k*-mer size, GenomeScope2 [66] was used to estimate heterozygosity, with the parameter for maximum *k*-mer coverage set to 900,000, as recommended by Mgwatyu et al. [67].

While visualizing the distribution of SNVs across scaffolds, we identified regions with loss of heterozygosity (LoH). To define these regions, we employed the LoH Caller tool [68]. We quantified the number of LoH events and calculated the coverage of LoH regions across the genomes of the 13 isolates. LoH events were categorized into two types: short LoH (less than 10 kb) and long LoH (greater than 100 kb). Additionally, the percentage of reduced heterozygosity level caused by LoH was determined by calculating the proportion of nucleotides that transitioned to a homozygous state in each isolate genome. The distribution of LoH regions of each isolate was visualized on the genome scaffolds using Python scripts. A matrix of LoH regions across isolates was constructed to generate a hierarchical clustering dendrogram and a heatmap displaying pairwise LoH overlaps among isolates, using a home-made Python scripts (https://github.com/PhanNgan/POPULATION-DIVERSITY). To evaluate the mode of natural selection acting on protein-coding genes affected by LoH, the ratio of non-synonymous to synonymous substitutions (dN/dS) was calculated for all protein-coding genes. Then, we compared this ratio for genes within LoH regions vs. the rest of the genes in the genome for each isolate. Besides, the efficacy of purifying selection (π_n_/π_s_) was also calculated and compared for genes in LoH and the rest of the genes.

### 5.5. Linkage disequilibrium and 4-gamete test using SNVs in the whole genome of the 13 *Mg* isolates

Homologous recombination in the *M. graminicola* genome can be revealed by studying the linkage disequilibrium between SNV markers as well as the proportion of pairs of markers passing the 4-gamete test [69], as a function of their distance along the genome. The SNV matrix between isolates was thus used for applying the 4-gamete test and estimating the linkage disequilibrium (LD) between nucleotide variants, using the script developed by Koutsovoulos et al. [70]. Note that SNVs in each isolate were phased to two haplotypes using WhatsHap [71] before inputting to 4-gamete test and LD. If a recombination event occurred between two bi-allelic sites (or markers) during meiosis I, four gametes will be formed at the end of meiosis II, including the two parental haplotypes and two recombinant haplotypes. Among the 13 *Mg* isolates, the number of haplotypes formed by two markers (at diploid state) on the same scaffold was counted. Then, the number of marker pairs was grouped according to their physical distances. At each inter-marker physical distance, the proportion of pairs of markers corresponding to the four products of meiosis (haplotypes) was calculated. The larger the distance between two markers is, the more likely recombination is expected to occur, so if meiotic recombination occurs, the proportion of marker pairs that pass the 4-gamete test should increase with the distance between markers.

The LD (r^2^) between the pairs of markers was calculated according to the following equation: r^2^ = (q_1_ q_2_ – p_1_ p_2_)^2^/r _1_r_2_ r_3_ r_4_, where q_1_ and q_2_ are the frequencies of two parental haplotypes (AB and ab); p_1_ and p_2_ are the frequencies of two recombinant haplotypes (Ab, aB); r_1,_ r_2,_ r_3_ and r_4_ are frequencies of four alleles (A, B, a, b) [72]. When the two markers are close one to another, there is less chance of recombination, resulting in a low frequency of recombinant haplotypes and a high r^2^ value. On the other hand, the greater distance between markers, the greater the possibility of recombination events leading to a decrease in the r^2^ value. The value of r^2^ can vary between 1 [no recombination, p_1_ p_2_ = 0, (q_1_ q_2_)^2^ = r_1_ r_2_ r_3_ r_4_] and 0 (when there is no more linkage between the parental variants due to frequent recombination, q_1_q_2_ = p_1_p_2_).

### 5.6. Principal component analysis using SNVs at the whole genome level

To cluster *M. graminicola* populations according to the identified SNVs, a Principal Component Analysis (PCA) was conducted. The SNV matrix between the 13 *Mg* isolates was used to perform a PCA using the SNPRelate package with default parameters [73]. We also calculated the fixation index (*F*_ST_; [74]) between the clusters using vcftools [62].

Bioinformatic tools automatically detected 44,527 SNVs across the whole genome, including both SNV tracts (LoH) and fixed SNVs caused by single-nucleotide mutations. However, LoH can interfere with linkage disequilibrium (LD) analysis by artificially inflating LD due to extended homozygous tracts. They can also affect the 4-gamete test by either increasing or reducing the ability to detect all four possible haplotypes (AB, Ab, aB, ab). Since current bioinformatic tools do not allow us to selectively filter single-nucleotide mutations within or near LoH regions at the whole-genome level, we opted to retain all 44,527 SNVs for recombination testing and principal component analysis (PCA). In parallel, we carefully selected a subset of fixed SNVs for LD analysis, the 4-gamete test, and PCA to refine and complement the broader genome-wide SNV analysis (See Supplementary Method 1).

### 5.7 Detection of copy number variants on the scaffolds of 13 *Mg* isolates

The cn.MOPS algorithm v1.24.0 [75] was applied to identify putative copy number variants (CNVs) in the genome sequence of the 13 isolates. First, the alignment of Illumina reads of each isolate against the haploid reference genome was used to calculate read coverage per 1- kb sliding window using Bedtools *multicov* [76]. Then a matrix of average read-coverage per 1-kb sliding window along the scaffolds of all 13 isolates was used as input to the cn.MOPS program. This program was run using medium normalization mode, and default values for all other parameters. cn.MOPS is a multiple sample read depth method that applies a Bayesian approach to decompose read coverage variations across multiple samples at each genomic position into integer copy numbers. By using Poisson distributions, noise was detected and removed, thus, reducing false positives. Finally, low complexity regions of the genome were excluded from this analysis. A matrix of copy number variation regions (CNVRs) across the genomes of 13 isolates, was created. Genome regions with a copy number of 2 were considered to represent the normal diploid state. Regions with copy numbers of CN0 and CN1 were interpreted as deletions and partial deletions, respectively. Regions with copy numbers greater than 2 (CN3–CN6) indicated duplications compared to the normal diploid state. The distribution of CNVRs (including duplication regions and deletion regions) was co-visualized with LoH regions on the genome scaffolds using Python scripts. Protein-coding genes overlapping CNV regions on at least 70% of their length were selected and considered involved in CNV. To assess the selection pressure on the coding genes affected by CNVs, the ratio of non-synonymous to synonymous substitutions (dN/dS) and the efficacy of purifying selection (π_n_ /π_s_) was also calculated using the SNVs located on these genes among isolates.

To further support the reliability of genome reorganization events (LoH and CNVs) in the Mg genome, 18 low-copy contigs were randomly selected and manually phased into two haplotypes (see Supplementary Method 2). Average read coverage analysis of their pairs of haplotypes (354 kb) would provide practical evidence of LoH and CNVs in the Mg genome (see Supplementary Method 2).

## Supporting information

Supplementary Result 1

Supplementary Result 2

## AUTHOR CONTRIBUTIONS

NTP conducted the bioinformatics analyses and drafted the original manuscript. EGJD contributed to the study’s conceptualization, prediction of secreted proteins and revision of the manuscript. GDK developed the bioinformatics pipelines for LD and four-gamete tests. CR executed gene annotation and wrote the corresponding methods and results. MLV participated in data analyses and improved the draft’s English language. GB and SB both contributed to the study’s conceptualization, funding acquisition, and revision of the manuscript.

## ACKNOWLEDGEMENT

This research was funded by the Consultative Group for International Agricultural Research (CGIAR) Program on Rice Agri-food Systems (CRP-RICE, 2017–2022) and the Research Institute for Development (IRD). Ngan Thi Phan was funded through Labex AGRO 2011- LABX-002, project no. 2002-010, (under I-Site Muse framework) coordinated by Agropolis Fondation. GB is a member of the CRBE laboratory supported by the French Laboratory of Excellence (LabEx) projects CEBA (ANR-10-LABX-25-01) and TULIP (ANR-10-LABX-41), managed by the French ANR. The authors want to thank Jamel Aribi (IRD-PHIM, France) for maintaining the nematode populations in greenhouse, Ndomassi Tando (IRD- DIADE, France) and the IRD itrop “Plantes Santé” bioinformatics platform for providing HPC resources and support for our research project.

## CONFLICT OF INTEREST STATEMENT

The authors declare no conflict of interest.

## DATA AVAILABILITY STATEMENT

The updated gene annotation for the *M. graminicola* genome described in this paper is available under NCBI BioProject xx, accession xx. Detailed procedural information regarding the analysis presented can be found in the GitHub repository at https://github.com/PhanNgan/POPULATION-DIVERSITY

## Notes

### Competing Interest Statement

The authors have declared no competing interest.

## REFERENCES

1. FAOSTAT. 2023. Available from: http://www.fao.org/faostat/en/#data/QC

2. Padgham JL, Duxbury JM, Mazid AM, Abawi GS, Hossain M. Yield loss caused by *Meloidogyne graminicola* on lowland rainfed rice in Bangladesh. J Nematol. 2004;36(1):42–8.

4. Bridge J, Luc M, Plowright RA. Nematode parasites of rice. In: Plant-parasitic nematodes in subtropical and tropical agriculture. CAB International; 2009. p. 69–108.

5. EPPO Global Database. 2023. Available from: https://gd.eppo.int/

6. Fanelli E, Cotroneo A, Carisio L, Troccoli A, Grosso S, Boero C, et al. Detection and molecular characterization of the rice root-knot nematode *Meloidogyne graminicola* in Italy. Eur J Plant Pathol. 2017;149(2):467–76. doi: 10.1007/s10658-017-1196-7

7. Golden AM, Birchfield W. *Meloidogyne graminicola* (Heteroderidae) a new species of root-knot nematode from grass. Proc Helminthol Soc Wash. 1965;32(2):228–31. Available from: https://www.cabdirect.org/cabdirect/abstract/19660801518

8. Golden AM, Birchfield W. Rice root-knot nematode (*Meloidogyne graminicola*) as a new pest of rice. Plant Dis Rep. 1968;52(6):243. Available from: https://www.cabdirect.org/cabdirect/abstract/19690806120

9. Monteiro AR, Ferraz LCCB. Encontro de *Meloidogyne graminicola* e primeiro ensaio de hospedabilidade no Brasil. Nematol Bras. 1988;12:149–50.

10. Rusinque L, Maleita C, Abrantes I, Palomares-Rius JE, Inácio ML. *Meloidogyne graminicola*—A threat to rice production: Review update on distribution, biology, identification, and management. Biology. 2021;10:1163. doi: 10.3390/biology10111163

11. Chapuis E, Besnard G, Andrianasetra S, Rakotomalala M, Nguyen HT, Bellafiore S. First report of the root-knot nematode *Meloidogyne graminicola* in Madagascar rice fields. Australas Plant Dis Notes. 2016;11(1):32. doi: 10.1007/s13314-016-0222-5

12. Bellafiore S, Jougla C, Chapuis É, Besnard G, Suong MN, Vu PN, et al. Intraspecific variability of the facultative meiotic parthenogenetic root-knot nematode (*Meloidogyne graminicola*) from rice fields in Vietnam. C R Biol. 2015;338(7):471–83. doi: 10.1016/j.crvi.2015.04.002

13. Phan NT, De Waele D, Lorieux M, Xiong L, Bellafiore S. A hypersensitivity-like response to *Meloidogyne graminicola* in rice (*Oryza sativa*). Phytopathology. 2018;108(4):521–8. doi: 10.1094/PHYTO-07-17-0235-R

14. De Waele D, Elsen A. Challenges in tropical plant nematology. Annu Rev Phytopathol. 2007;45(1):457–85. doi: 10.1146/annurev.phyto.45.062806.094438

15. Besnard G, Thi-Phan N, Ho-Bich H, Dereeper A, Nguyen HT, Quénéhervé P, et al. On the close relatedness of two rice-parasitic root-knot nematode species and the recent expansion of *Meloidogyne graminicola* in Southeast Asia. Genes (Basel). 2019;10(2). doi: 10.3390/genes10020175

16. Mondal S, Purohit A, Hazra A, Sharma R, Gupta P, Banerjee T, et al. Intraspecific variability of rice root knot nematodes across diverse agroecosystems for sustainable management. Sci Rep. 2024;14:30032. doi: 10.1038/s41598-024-73980-x.

17. Carneiro RMDG, Almeida MR, Quénéhervé P. Enzyme phenotypes of *Meloidogyne* spp. populations. Nematology. 2000;2:645–54. https://www.cabi.org/isc/abstract/20013029216

18. Pokharel RR, Abawi GS, Duxbury JM, Smat CD, Wang X, Roberts PA. Variability and the recognition of two races in *Meloidogyne graminicola*. Australas Plant Pathol. 2010;39(4):326–33. doi: 10.1071/AP09100

19. Gibson AK, Delph LF, Lively CM. The two-fold cost of sex: Experimental evidence from a natural system. Evol Lett. 2017;1(1):6–15. doi: 10.1002/evl3.1

20. Burt A. Perspective: Sex, recombination, and the efficacy of selection—was Weismann right? Evolution. 2000;54(2):337–51. doi: 10.1111/j.0014-3820.2000.tb00038.x

21. Kondrashov AS. Selection against harmful mutations in large sexual and asexual populations. Genet Res. 1982;40(3):325–32.

22. Lynch M, Bürger R, Butcher D, Gabriel W. The mutational meltdown in asexual populations. J Hered. 1993;84(5):339–44. doi: 10.1093/oxfordjournals.jhered.a111354

23. Glémin S, François CM, Galtier N. Genome evolution in outcrossing vs. selfing vs. asexual species. In: Anisimova M, editor. Evolutionary Genomics: Statistical and Computational Methods. Springer; 2019. p. 331–69. doi: 10.1007/978-1-4939-9074-0_11

24. Maynard Smith J. Evolution: Contemplating life without sex. Nature. 1986;324(6095):300–1. doi: 10.1038/324300a0

25. Archetti M. Recombination and loss of complementation: A more than two-fold cost for parthenogenesis. J Evol Biol. 2004;17(5):1084–97. doi: 10.1111/j.1420-9101.2004.00745.x

26. Vakhrusheva OA, Mnatsakanova EA, Galimov YR, Neretina TV, Gerasimov ES, Ozerova SG, et al. Recombination in a natural population of the bdelloid rotifer *Adineta vaga*. bioRxiv. 2018;489393. doi: 10.1101/489393

27. Jaron KS, Bast J, Nowell RW, Ranallo-Benavidez TR, Robinson-Rechavi M, Schwander T. Genomic features of parthenogenetic animals. J Hered. 2021;112(1):19–33. doi: 10.1093/jhered/esaa031

28. Hassanaly-Goulamhoussen R, De Carvalho AR, Marteu-Garello N, Péré A, Favery B, Da Rocha M, et al. Chromatin landscape dynamics in the early development of the plant parasitic nematode *Meloidogyne incognita*. Front Cell Dev Biol. 2021;9:765690. doi: 10.3389/fcell.2021.765690

28. Kozlowski DKL, Hassanaly-Goulamhoussen R, Da Rocha M, Koutsovoulos GD, Bailly-Bechet M, Danchin EGJ. Movements of transposable elements contribute to the genomic plasticity and species diversification in an asexually reproducing nematode pest. Evol Appl. 2021;14(7):1844–66. doi: 10.1111/eva.13246

29. Tralamazza SM, Gluck-Thaler E, Feurtey A, Croll D. Copy number variation introduced by a massive mobile element facilitates global thermal adaptation in a fungal wheat pathogen. Nat Commun. 2024;15:5728. doi: 10.1038/s41467-024-49913-7

30. Castagnone-Sereno P, Mulet K, Danchin EGJ, Koutsovoulos GD, Karaulic M, Rocha MD, et al. Gene copy number variations as signatures of adaptive evolution in the parthenogenetic, plant-parasitic nematode *Meloidogyne incognita*. Mol Ecol. 2019;28(10):2559–72. doi: 10.1111/mec.15095

31. Triantaphyllou AC. Gametogenesis and the chromosomes of two root-knot nematodes, Meloidogyne graminicola and M. naasi. J Nematol. 1969;1(1):62–71. pmid: 19325656

32. McDonald BA, Linde C. Pathogen population genetics, evolutionary potential, and durable resistance. Annu Rev Phytopathol. 2002;40(1):349–79. doi: 10.1146/annurev.phyto.40.120501.101443

33. Phan NT, Orjuela J, Danchin EGJ, Klopp C, Perfus-Barbeoch L, Kozlowski DK et al. Genome structure and content of the rice root-knot nematode (*Meloidogyne graminicola*). Ecol Evol. 2020;10(20):11006–21. doi: 10.1002/ece3.6680

34. Somvanshi VS, Dash M, Bhat CG, Budhwar R, Godwin J, Shukla RN, Patrignani A, Schlapbach R, Rao U. An improved draft genome assembly of *Meloidogyne graminicola* IARI strain using long-read sequencing. Gene. 2021;793:145748. doi: 10.1016/j.gene.2021.145748

35. Dai D, Xie C, Zhou Y, Bo D, Zhang S, Mao S, et al. Unzipped chromosome-level genomes reveal allopolyploid nematode origin pattern as unreduced gamete hybridization. Nat Commun. 2023;14:7156. doi: 10.1038/s41467-023-42700-w

36. Besnard G, Jühling F, Chapuis E, Zedane L, Lhuillier E, Mateille T, et al. Fast assembly of the mitochondrial genome of a plant parasitic nematode (*Meloidogyne graminicola*) using next-generation sequencing. C R Biol. 2014;337:295–301. doi: 10.1016/j.crvi.2014.03.003

37. Martin M. Cutadapt removes adapter sequences from high-throughput sequencing reads. EMBnet J. 2011;17(1):10–2. doi: 10.14806/ej.17.1.200

38. Jiang H, Lei R, Ding S-W, Zhu S. Skewer: A fast and accurate adapter trimmer for next-generation sequencing paired-end reads. BMC Bioinformatics. 2014;15(1):182. doi: 10.1186/1471-2105-15-182

39. Andrews S. FastQC: A quality control tool for high throughput sequence data. 2010. Available from: http://www.bioinformatics.babraham.ac.uk/projects/fastqc/

40. Kumar S, Jones M, Koutsovoulos G, Clarke M, Blaxter M. Blobology: Exploring raw genome data for contaminants, symbionts and parasites using taxon-annotated GC-coverage plots. Front Genet. 2013;4(237):237. doi: 10.3389/fgene.2013.00237

41. Bankevich A, Nurk S, Antipov D, Gurevich AA, Dvorkin M, Kulikov AS, et al. SPAdes: A new genome assembly algorithm and its applications to single-cell sequencing. J Comput Biol. 2012;19(5):455–77. doi: 10.1089/cmb.2012.0021

42. Laetsch DR, Blaxter ML. BlobTools: Interrogation of genome assemblies. F1000Res. 2017;6:1287. doi: 10.12688/f1000research.12232.1

43. Langmead B, Salzberg SL. Fast gapped-read alignment with Bowtie 2. Nat Methods. 2012;9(4):357–9. doi: 10.1038/nmeth.1923

44. Li H. lh3/seqtk: Toolkit for processing sequences in FASTA/Q formats [C]. 2020. Available from: https://github.com/lh3/seqtk (Original work published 2012)

45. Sallet E, Gouzy J, Schiex T. EuGene: An automated integrative gene finder for eukaryotes and prokaryotes. Methods Mol Biol. 2019;1962:97–120. doi:10.1007/978-1-4939-8958-4_6

46. Somvanshi VS, Tathode M, Shukla RN, Rao U. An improved draft genome assembly of *Meloidogyne graminicola* IARI strain using long-read sequencing. Gene. 2021 Aug;793:145748. doi: 10.1016/j.gene.2021.145748.

47. Howe KL, Bolt BJ, Shafie M, Kersey P, Berriman M. WormBase ParaSite - a comprehensive resource for helminth genomics. Mol Biochem Parasitol. 2017;215:2–10. doi:10.1016/j.molbiopara.2016.11.005

48. UniProt Consortium T. UniProt: The universal protein knowledgebase. Nucleic Acids Res. 2018;46:2699–2699. doi:10.1093/nar/gky092

49. Bao W, Kojima KK, Kohany O. Repbase update, a database of repetitive elements in eukaryotic genomes. Mob DNA. 2015;6:11. doi:10.1186/s13100-015-0041-9

50. Haas BJ, Papanicolaou A, Yassour M, Grabherr M, Blood PD, Bowden J, et al. De novo transcript sequence reconstruction from RNA-seq using the Trinity platform for reference generation and analysis. Nat Protoc. 2013;8:1494–512. doi:10.1038/nprot.2013.084

51. Pertea M, Pertea GM, Antonescu CM, Chang TC, Mendell JT, Salzberg SL et al. StringTie enables improved reconstruction of a transcriptome from RNA-seq reads. Nat Biotechnol. 2015;33:290–5. doi:10.1038/nbt.3122

52. Schulz MH, Zerbino DR, Vingron M, Birney E. Oases: Robust de novo RNA-seq assembly across the dynamic range of expression levels. Bioinformatics. 2012;28:1086–1092. doi:10.1093/bioinformatics/bts094

53. Simão FA, Waterhouse RM, Ioannidis P, Kriventseva EV, Zdobnov EM. BUSCO: Assessing genome assembly and annotation completeness with single-copy orthologs. Bioinformatics.

54. Almagro Armenteros JJ, Tsirigos KD, Sønderby CK, Petersen TN, Winther O, Brunak S, et al. SignalP 5.0 improves signal peptide predictions using deep neural networks. Nat Biotechnol. 2019;37:420–3. doi: 10.1038/s41587-019-0036-z

55. Krogh A, Larsson B, von Heijne G, Sonnhammer ELL. Predicting transmembrane protein topology with a hidden Markov model: application to complete genomes. J Mol Biol. 2001;305:567–80. doi: 10.1006/jmbi.2000.4315

56. Jones P, Binns D, Chang H-Y, Fraser M, Li W, McAnulla C, et al. InterProScan 5: genome-scale protein function classification. Bioinformatics. 2014;30:1236–40. doi: 10.1093/bioinformatics/btu031

57. Mitchell AL, Attwood TK, Babbitt PC, Blum M, Bork P, Bridge A, et al. InterPro in 2019: Improving coverage, classification and access to protein sequence annotations. Nucleic Acids Res. 2019;47(D1):D351–60. doi: 10.1093/nar/gky1100

58. Li H, Durbin R. Fast and accurate short read alignment with Burrows-Wheeler transform. Bioinformatics. 2009;25(14):1754–60. doi: 10.1093/bioinformatics/btp324

59. Li H, Handsaker B, Wysoker A, Fennell T, Ruan J, Homer N, et al. The sequence alignment/Map format and SAMtools. Bioinformatics. 2009;25(16):2078–9. doi: 10.1093/bioinformatics/btp352

60. McKenna A, Hanna M, Banks E, Sivachenko A, Cibulskis K, Kernytsky A, et al. The Genome Analysis Toolkit: A MapReduce framework for analyzing next-generation DNA sequencing data. Genome Res. 2010;20(9):1297–303. doi: 10.1101/gr.107524.110

61. Garrison E, Marth G. Haplotype-based variant detection from short-read sequencing. ArXiv:1207.3907 [q-Bio]. 2012. doi: 10.48550/arXiv.1207.3907

62. Danecek P, Auton A, Abecasis G, Albers CA, Banks E, DePristo MA, et al. The variant call format and VCFtools. Bioinformatics. 2011;27(15):2156–8. doi: 10.1093/bioinformatics/btr330

63. Cingolani P, Platts A, Wang LL, Coon M, Nguyen T, Wang L, et al. A program for annotating and predicting the effects of single nucleotide polymorphisms, SnpEff: SNPs in the genome of Drosophila melanogaster strain w1118; iso-2; iso-3. Landes Biosci. 2012;6(2):80-92. doi: 10.4161/fly.19695

64. Guéguen L, Gaillard S, Boussau B, Gouy M, Groussin M, Rochette NC, et al. Bio++: efficient extensible libraries and tools for computational molecular evolution. Mol Biol Evol. 2013;30:1745–50. doi: 10.1093/molbev/mst097

65. Marçais G, Kingsford C. A fast, lock-free approach for efficient parallel counting of occurrences of k-mers. Bioinformatics. 2021;27(6):764–70. doi: 10.1093/bioinformatics/btr011

66. Ranallo-Benavidez TR, Jaron KS, Schatz MC. GenomeScope 2.0 and Smudgeplot for reference-free profiling of polyploid genomes. Nat Commun. 2020;11:1432. doi: 10.1038/s41467-020-14998-3

67. Mgwatyu Y, Stander AA, Ferreira S, Williams W, Hesse U. Rooibos (*Aspalathus linearis*) genome size estimation using flow cytometry and k□mer analyses. Plants. 2020;9(2):270. doi: 10.3390/plants9020270

68. Georgeson, Peter. supernifty/LOHdeTerminator. 2018. 19 décembre 2024. GitHub, https://github.com/supernifty/LOHdeTerminator.

69. Hudson RK. Statistical properties of the number of recombination events in the history of a sample of DNA sequences. Genetics. 1985;111(1):147–64.

70. Koutsovoulos GD, Marques E, Arguel M-J, Duret L, Machado ACZ, Carneiro RMDG, et al. Population genomics supports clonal reproduction and multiple gains and losses of parasitic abilities in the most devastating nematode plant pest. Evol Appl. 2019;13(2):442–57. doi: 10.1111/eva.12881

71. Martin M, Patterson M, Garg S, Fischer SO, Pisanti N, Klau GW, et al. WhatsHap: Fast and accurate read-based phasing. bioRxiv. 2016;085050. doi: 10.1101/085050

72. Slatkin M. Linkage disequilibrium—understanding the evolutionary past and mapping the medical future. Nat Rev Genet. 2008;9:477–485. doi: 10.1038/nrg2361

73. Zheng X, Levine D, Shen J, Gogarten SM, Laurie C, Weir BS. A high-performance computing toolset for relatedness and principal component analysis of SNP data. Bioinformatics. 2012;28(24):3326–3328. doi: 10.1093/bioinformatics/bts606

74. Weir BS, Cockerham CC. Estimating *F*-statistics for the analysis of population structure.

75. Klambauer G, Schwarzbauer K, Mayr A, Clevert D-A, Mitterecker A, Bodenhofer U, et al. cn.MOPS: Mixture of Poissons for discovering copy number variations in next-generation sequencing data with a low false discovery rate. Nucleic Acids Res. 2012;40(9):e69. doi: 10.1093/nar/gks003

76. Quinlan AR, Hall IM. BEDTools: A flexible suite of utilities for comparing genomic features. Bioinformatics. 2010;26(6):841–2. doi: 10.1093/bioinformatics/btq033

77. Mota APZ, Koutsovoulos GD, Perfus-Barbeoch L, Despot-Slade E, Labadie K, Aury JM, et al. Unzipped genome assemblies of polyploid root-knot nematodes reveal unusual and clade-specific telomeric repeats. Nat Commun. 2024;15:773. doi: 10.1038/s41467-024-44914-y

78. Rehman S, Gupta VK, Goyal AK. Identification and functional analysis of secreted effectors from phytoparasitic nematodes. BMC Microbiol. 2016;16:48. doi: 10.1186/s12866-016-0632-8

79. Romiguier J, Gayral P, Ballenghien M, Bernard A, Cahais V, Chenuil A, et al. Comparative population genomics in animals uncovers the determinants of genetic diversity. Nature. 2014;515:261–3. doi: 10.1038/nature13685

80. Lewin B. Genes IV. Oxford: Oxford University Press; 1990.

81. Lancaster SM, Payen C, Heil CS, Dunham MJ. Fitness benefits of loss of heterozygosity in *Saccharomyces* hybrids. Proc Natl Acad Sci U S A. 2020;117(42):26669–77. doi: 10.1073/pnas.2018633117

82. Heil SC. Loss of heterozygosity and its importance in evolution. J Mol Evol. 2023;91(3):369–77. doi: 10.1007/s00239-022-10088-8

83. Leffler EM, Bullaughey K, Matute DR, Meyer WK, Ségurel L, Venkat A, et al. Revisiting an old riddle: What determines genetic diversity levels within species? PLoS Biol. 2012;10(9):e1001388. doi: 10.1371/journal.pbio.1001388

84. Lacy KD, Hart T, Kronauer DJC. Co-inheritance of recombined chromatids maintains heterozygosity in a parthenogenetic ant. Nat Ecol Evol. 2024;8:1522–33. doi: 10.1038/s41559-024-02455-z

85. Liu QL, Thomas VP, Williamson VM. Meiotic parthenogenesis in a root-knot nematode results in rapid genomic homozygosity. Genetics. 2007;176:1483–90. doi: 10.1534/genetics.107.071134

86. Archetti M. Complementation, genetic conflict, and the evolution of sex and recombination. J Hered. 2010;101:21–33. doi: 10.1093/jhered/esq009

87. Bi K, Bogart JP. Time and time again: Unisexual salamanders (genus *Ambystoma*) are the oldest unisexual vertebrates. BMC Evol Biol. 2010;10:238. doi: 10.1186/1471-2148-10-238

88. Hojsgaard D, Hörandl E. A little bit of sex matters for genome evolution in asexual plants. Front Plant Sci. 2015;6:82. doi: 10.3389/fpls.2015.00082

89. Lunt DH, Kumar S, Koutsovoulos G, Blaxter ML. The complex hybrid origins of the root knot nematodes revealed through comparative genomics. PeerJ. 2014;2:e356.

90. Forche A, Alby K, Schaefer D, Johnson AD, Berman J, Bennett RJ. The parasexual cycle in *Candida albicans* provides an alternative pathway to meiosis for the formation of recombinant strains. PLoS Biol. 2008;6:e110. doi: 10.1371/journal.pbio.0060110

91. Schön I, Martens K, van Dijk P. Lost sex: The evolutionary biology of parthenogenesis. Berlin (Germany): Springer; 2009.

92. Engelstädter J. Asexual but not clonal: evolutionary processes in automictic populations. Genetics. 2017;206:993–109. doi: 10.1534/genetics.116.196873

93. Flynn JM, Chain FJJ, Schoen DJ, Cristescu ME. Spontaneous mutation accumulation in *Daphnia pulex* in selection-free vs. competitive environments. Mol Biol Evol. 2017;34(1):160–73. doi: 10.1093/molbev/msw234

94. Ford CB, Funt JM, Abbey D, Issi L, Guiducci C, Martinez DA, et al. The evolution of drug resistance in clinical isolates of *Candida albicans*. eLife. 2015;4:e00662. doi: 10.7554/eLife.00662

95. Flot JF, Hespeels B, Li X, Noel B, Arkhipova I, Danchin EGJ, et al. Genomic evidence for ameiotic evolution in the bdelloid rotifer *Adineta vaga*. Nature. 2013;500(7463):453–7. doi: 10.1038/nature12326

96. Latta LC, Tucker KN, Haney RA. The relationship between oxidative stress, reproduction, and survival in a bdelloid rotifer. BMC Ecol. 2019;19(1):7. doi: 10.1186/s12898-019-0223-2

97. Hespeels B, Penninckx S, Cornet V, Bruneau L, Bopp C, Baumlé V, et al. Iron ladies – How desiccated asexual rotifer *Adineta vaga* deal with X-rays and heavy ions? Front Microbiol. 2020;11:1792. doi: 10.3389/fmicb.2020.01792

98. Haegeman A, Mantelin S, Jones JT, Gheysen G. Functional roles of effectors of plant-parasitic nematodes. Gene. 2012;492(1):19–31. doi: 10.1016/j.gene.2011.10.040

99. Chen J, Glémin S, Lascoux M. Genetic diversity and the efficacy of purifying selection across plant and animal species. Mol Biol Evol. 2017;34(6):1417–28. doi: 10.1093/molbev/msx088

100. Schughart K, Kappen C, Ruddle FH. Duplication of large genomic regions during the evolution of vertebrate homeobox genes. Proc Natl Acad Sci U S A. 1989;86(18):7067–7071. doi: 10.1073/pnas.86.18.7067

101. Alkan C, Coe BP, Eichler EE. Genome structural variation discovery and genotyping. Nat Rev Genet. 2011;12:363–76. doi: 10.1038/nrg2958

102. Arlt MF, Rajendran S, Birkeland SR, Wilson TE, Glover TW. Copy number variants are produced in response to low-dose ionizing radiation in cultured cells. Environ Mol Mutagen. 2014;55(2):103–13. doi: 10.1002/em.21840

103. Bussotti G, Gouzelou E, Boité MC, Kherachi I, Harrat Z, Eddaikra N, et al. *Leishmania* genome dynamics during environmental adaptation reveal strain-specific differences in gene copy number variation, karyotype instability, and telomeric amplification. MBio. 2018;9(6). doi: 10.1128/mBio.01399-18

104. Dasmeh P, Serohijos AW, Kepp KP, Shakhnovich EI. The influence of selection for protein stability on dN/dS estimations. Genome Biol Evol. 2014;6(10):2956–67. doi: 10.1093/gbe/evu231.

105. Buckingham LJ, Ashby B. Coevolutionary theory of hosts and parasites. J Evol Biol. 2022;35(2):205–24. doi: 10.1111/jeb.13981

106. Cook DE, Lee TG, Guo X, Melito S, Wang K, Bayless AM, et al. Copy number variation of multiple genes at Rhg1 mediates nematode resistance in soybean. Science. 2012;338(6111):1206–9. doi: 10.1126/science.1228746

107. Patil GB, Lakhssassi N, Wan J, Song L, Zhou Z, Klepadlo M, et al. Whole□genome re□sequencing reveals the impact of the interaction of copy number variants of the rhg1 and Rhg4 genes on broad□based resistance to soybean cyst nematode. Plant Biotechnol J. 2019;17(8):1595– 611. doi: 10.1111/pbi.13086

108. Nguyen HT, Vang S, Phan NT, Czernic P, Trinh PQ, Ha CV, et al. Identification and characterization of a virulent population of *Meloidogyne graminicola*. Australas Plant Pathol. 2023;52:391–405. doi: 10.1007/s13313-023-00926-8

109. Alberts B, Johnson A, Lewis J, Raff M, Roberts K, Walter P. General recombination. In: Molecular Biology of the Cell. 4th ed. 2002. Available from: https://www.ncbi.nlm.nih.gov/books/NBK26898/

110. Denver DR, Morris K, Lynch M, Thomas WK. High mutation rate and predominance of insertions in the *Caenorhabditis elegans* nuclear genome. Nature. 2004;430(7000):679–82. doi: 10.1038/nature02697

111. Neiman M, Taylor DR. The causes of mutation accumulation in mitochondrial genomes. Proc R Soc B Biol Sci. 2009;276(1660):1201–9. doi: 10.1098/rspb.2008.1758

112. Goodwin S, McPherson JD, McCombie WR. Coming of age: ten years of next-generation sequencing technologies. Nat Rev Genet. 2011;12(10):671–82. doi: 10.1038/nrg3025

113. Nei M, Gojobori T. Simple methods for estimating the numbers of synonymous and nonsynonymous nucleotide substitutions. Mol Biol Evol. 1986;3:418–26. doi: 10.1093/oxfordjournals.molbev.a040410

114. Lynch M. The Origins of Genome Architecture. Sunderland (MA): Sinauer; 2007.

115. Vranken S, Scheben A, Batley J, Wernberg T, Coleman MA. Genomic consequences and selection efficacy in sympatric sexual versus asexual kelps. Front Mar Sci. 2022;9:921912. doi: 10.3389/fmars.2022.921912

116. Liu Q, Weissman DB. Population structure can reduce clonal interference when sexual reproduction and dispersal are synchronized. bioRxiv. 2023. doi: 10.1101/2023.07.10.548343

