## Supplementary Method 1 for "Genomic rearrangements promote diversification of a facultative meiotic parthenogenetic nematode pest (*Meloidogyne graminicola*)"

**Identification of a subset of single-nucleotide mutations among the 13 *Mg* isolates to further support recombination testing, PCA, and haplotype network reconstruction**

Visualization of positions and genotypes of nucleotide variants among the genome of 13 *Mg* isolates on scaffolds at section 2.2 show the clusters of homozygous SNVs (“1/1”) that represent genome reorganization events. Genomic regions showing suites of homozygous SNVs (“1/1”) separated by less than 100 bp were detected and categorized as genome reorganization events (LoH). Meanwhile, sporadically distributed SNVs (“1/1” and “2/2”) in the genome were classified as single nucleotide mutations.

Since current bioinformatic tools do not allow us to selectively filter single-nucleotide mutations within or near LoH regions at the whole-genome level, we manually selected (by checking on the alignment) a subset of single mutations which were 1) no showing heterozygous genotype (0/1, 0/2) in any among 13 isolates, 2) highly supported (minimum coverage of 10×); 3) not belonging to duplicate regions; and 4) not located near mononucleotide stretches (e.g. poly-T or poly-A) that may cause sequencing errors. Subsequently, this set of nucleotide variants was also used as input into the recombination test (4-gamete test and LD) and the principal component analysis as described above. Finally, this SNVs set was aligned to construct a median joining network using PopArt v1.7 [1].

REFERENCES

1. Leigh JW, Bryant D. popart: full-feature software for haplotype network construction. Meth Ecol Evol. 2015;6(9):1110–6. doi. 10.1111/2041-210X.12410
