## Supplementary Method 2 for "Genomic rearrangements promote diversification of a facultative meiotic parthenogenetic nematode pest (*Meloidogyne graminicola*)"

**Set of nuclear divergent copies for read coverage analysis and nucleotide variant calling to support CNVs, LoH, and potentially recombination**

The current assembly tools using a combination of both long and short reads were not able to assemble two distinct phased haplotypes for the *M. graminicola* genome. Aware of that, two-state genotypes were considered when selecting the nucleotide variants among isolates for principal component analysis and recombination test. Furthermore, average read coverage analysis of the two haplotypes could indicate the deletion or duplication of one or both alleles, thereby providing evidence of CNVs and LoH regions in the *Mg* genome.

We randomly targeted 18 low-copy contigs (177 kb; including TAA6 and ACC6 contigs initially assembled in [2]), manually phased into two haplotypes, creating 18 pairs (354 kb, 0.43% of the genome) of divergent contigs with sequence divergence ranging from 0.3 to 5.7% (Table S2). These 18 genomic regions were extracted from the reference consensus genome of *M. graminicola*. For each contig, two divergent copies were separately assembled using the methods described in Besnard et al. [2]. Briefly, the Illumina sequences of the Mg-VN18 isolate were mapped to all 18 nuclear contigs using the BWA-MEM software [3]. The mapping of paired-end reads on these contigs was also visualized and inspected with Geneious v6 [4]. Then, based on the linkage between paired-end reads, reads were carefully phased and concatenated into separate haplotypes, resulting in the assembly of two divergent contig pairs (i.e. two haplotypes). The Illumina reads of Mg-VN18 were again mapped to the assembled divergent copies of each contig to correct the sequence of the two copies. Finally, haplotypes of these contigs were separated in 36 sequences covering 354 kb, and used as a new reference.

The Illumina reads of the 13 isolates were then mapped to these 36 sequences using BWA-MEM. Each alignment was visualized on Geneious to manually verify that reads were clearly separated into two divergent copies with a correct read phasing. Then, the average read-coverage was calculated for each region. In addition, variants on the divergent copies were called among the isolates using the same method as described above. All detected variants were manually validated by a visual inspection of the alignment with Geneious. The quality control of the read mapping and variant search were also performed on five populations (Mg-Bali, Mg-Borneo, Mg-Java2, Mg-L2, and Mg-VN18) with very high sequencing coverage (288 to 568×).

REFERENCES

2. Besnard G, Thi-Phan N, Ho-Bich H, Dereeper A, Nguyen HT, Quénéhervé P, et al. On the close relatedness of two rice-parasitic root-knot nematode species and the recent expansion of *Meloidogyne graminicola* in Southeast Asia. Genes (Basel). 2019;10(2). doi: 10.3390/genes10020175

3. Li H, Durbin R. Fast and accurate short read alignment with Burrows-Wheeler transform. Bioinformatics. 2009;25(14):1754–60. doi: 10.1093/bioinformatics/btp324

4. Kearse M, Moir R, Wilson A, Stones-Havas S, Cheung M, Sturrock S, et al. Geneious basic: an integrated and extendable desktop software platform for the organization and analysis of sequence data. Bioinformatics. 2012;28:1647–9. doi: 10.1093/bioinformatics/bts199
