## Supplementary Result 1 for "Genomic rearrangements promote diversification of a facultative meiotic parthenogenetic nematode pest (*Meloidogyne graminicola*)"

**Few fixed mutations selected at whole genome level and on 18 manually curated contigs**

After careful filtration and visual inspection, only 40 fixed single nucleotide mutations (in total of 44,527 SNVs) were selected among 12 isolates in comparison to Mg-VN18 (See Supplementary Method 1). We excluded SNVs that were not associated with genome rearrangements, as well as sporadic SNVs where a heterozygous state occurred in any isolate. Therefore, the 40 selected SNVs do not represent the total diversity of the isolates but rather a subset of fixed SNVs. More point mutations were found in Mg-C25, Mg-P, Mg-C21, and Mg-VN27 isolates compared to Mg-VN18 genomic sequence (Figure S5A). The principal component analysis of 40 selected SNVs did not produce well-defined clusters possibly due to weak SNV signals (Figure S5B). However, it highlighted a close relationship among the eight isolates Mg-Bali, Mg-Borneo, Mg-C21, Mg-Java2, Mg-P, Mg-VN11, Mg-VN6, and Mg-L1 (Figure S5B).

Careful visual inspection of reads mapping on both haplotypes of the 18 manually curated genomic regions (354 kb) revealed only two SNVs on G859_T2 (one unique to Mg-C21 and another to Mg-P) and one InDel on G859_T1 (unique to Mg-C21) (See Supplementary Method 2, Figure S6). Therefore, these few mutations on the curated phased contigs did not allow us to re-run the principal component analysis and recombination test.
