## Supplementary Result 2 for "Genomic rearrangements promote diversification of a facultative meiotic parthenogenetic nematode pest (*Meloidogyne graminicola*)"

**Average read coverage analysis revealed the presence of CNVs and LoH on 18 manually curated contigs**

Visual inspection of reads mapping on the divergent haplotypes of 18 pairs of contigs (354 kb; 0.43% of the genome) revealed that some isolates have only one type (i.e. one haplotype is missing in some isolates). Note that reads assignment to each haplotype of all targeted regions was unambiguous, as shown on Figure S6. Therefore, for a given genomic contig, the average read-coverage on the two divergent haplotypes in each isolate showed their presence or absence. While all divergent haplotypes were present in four isolates (Mg-VN18, Mg-VN11, Mg-Brazil, and Mg-L2), some were absent in the remaining nine (Figure S7). The GB2_T2 contig was absent in eight of the 13 isolates (Mg-C21, Mg-C25, Mg-Vn27, Mg-Bali, Mg-Borneo, Mg-Java2, Mg-L1 and Mg-VN6), contig G315_T1 was absent in two isolates (Mg-L1 and Mg-C25), and finally, nine other contigs were absent in only one isolate (Figure S7). Out of these 21 cases (on 11 contigs) where a haplotype was missing, this was systematically accompanied by the doubling of sequencing coverage of its counterpart in 20 cases (on ten contigs). As an example, in Mg-C21, contig GB2_T2 was absent but its homologous GB2_T1 showed a 32× coverage, approximately twice the average sequencing depth of this isolate (15×; Figure S8). For these ten contigs, these observations suggest that one haplotype was present at the homozygous state representing loss of heterozygosity regions. Only the contig ACC4_T1 and its counterpart ACC4_T2 did not appear to follow this rule in the Mg-C25 isolate. In this case, absence of ACC4_T1 was not accompanied by higher read coverage of its counterpart ACC4_T2, suggesting a single deletion of ACC4_T1 leading to a hemizygous region (Figure S7).

As the sequencing coverage was not high for some isolates (ca. 15-30×), the analysis was repeated using five isolates including Mg-Bali, Mg-Borneo, Mg-L2, Mg-Java2, and Mg-VN18, for which high sequencing coverage is available (288-568×). The high read coverage in these five isolates allowed us to confirm five cases (out of more than 20) of LoH, characterized by sequencing coverage doubling of the extant contig at the expense of its absent counterpart (Figure S8). Additionally, it clearly revealed the deletion of G2728_T1 in the Mg-Bali isolate and the duplication of G1112-T1 in Mg-Borneo (Figure S8).
