## Supplementary Figure S1 for "Genomic rearrangements promote diversification of a facultative meiotic parthenogenetic nematode pest (*Meloidogyne graminicola*)"

### BUSCO completeness of four predicted proteome versions of *Meloidogyne graminicola* including ASM1477313v1 (10,331 proteins, [5]); Eugene_Mg (15,518 proteins, this study); PRJNA411966 (14,062 proteins, [6]), and T2T (12,968 proteins, [7]) using BUSCO version 5.5.0 and three different databases eu10 (eukaryota_odb10, n = 255), me10 (metazoa_odb10, n = 954); and ne10 (nematoda_odb10, n = 3131)

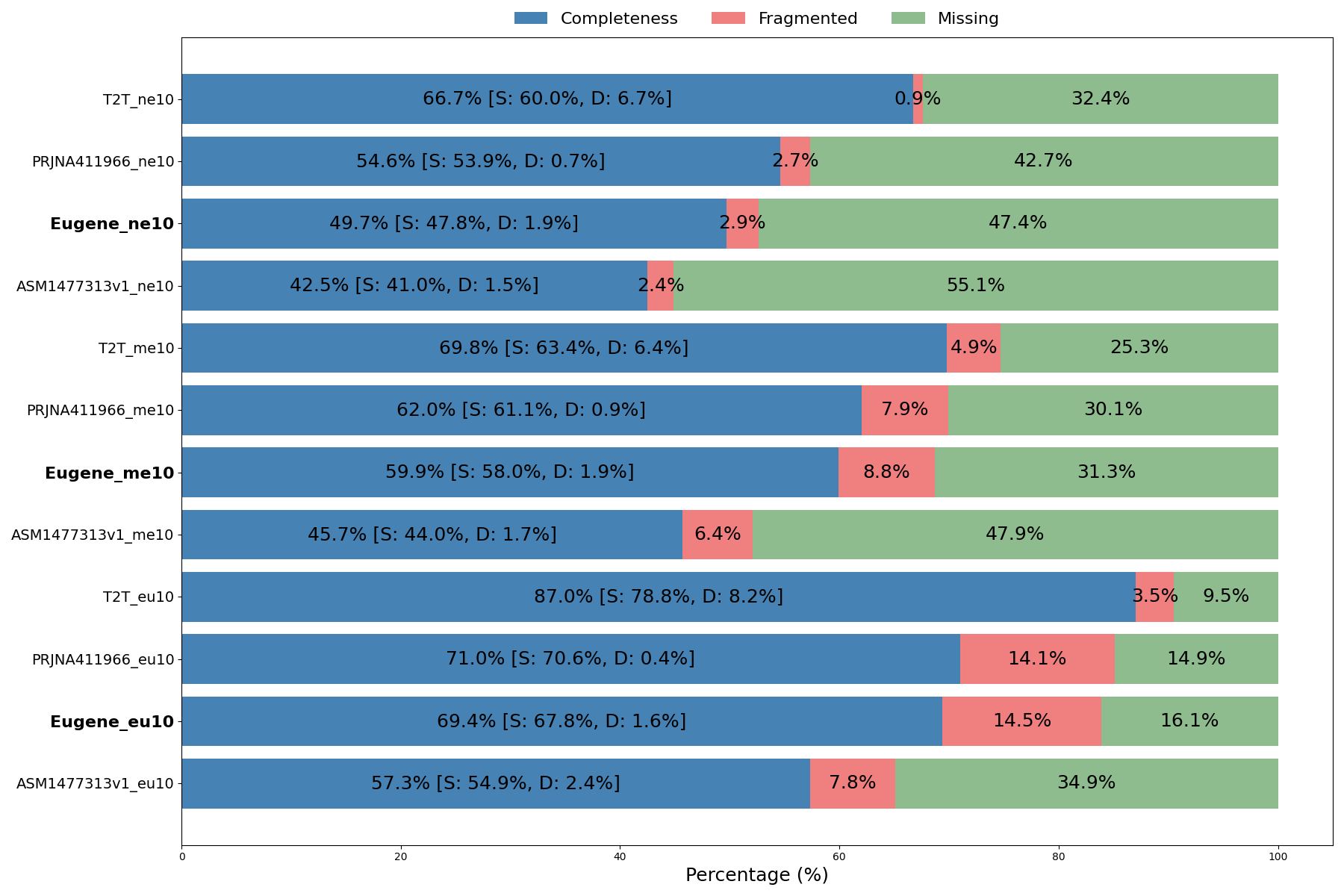

REFERENCES

5. Phan NT, Orjuela J, Danchin EGJ, Klopp C, Perfus-Barbeoch L, Kozlowski DK et al. Genome structure and content of the rice root-knot nematode (*Meloidogyne graminicola*). Ecol Evol. 2020;10(20):11006–21. doi: 10.1002/ece3.6680

6. Somvanshi VS, Dash M, Bhat CG, Budhwar R, Godwin J, Shukla RN, Patrignani A, Schlapbach R, Rao U. An improved draft genome assembly of *Meloidogyne graminicola* IARI strain using long-read sequencing. *Gene*. 2021;793:145748. doi: 10.1016/j.gene.2021.145748

7. Dai D, Xie C, Zhou Y, Bo D, Zhang S, Mao S, et al. Unzipped chromosome-level genomes reveal allopolyploid nematode origin pattern as unreduced gamete hybridization. Nat Commun. 2023;14:7156. doi: 10.1038/s41467-023-42700-w
