## Supplementary Figure S2A for "Genomic rearrangements promote diversification of a facultative meiotic parthenogenetic nematode pest (*Meloidogyne graminicola*)"

### Visualization of 693,013 SNVs, including both homozygous and heterozygous genotypes among 13 isolates across the 283 scaffolds of the reference genome.

The lengths of the scaffolds are presented to scale. Each disc represents a set of scaffolds ranked from the biggest to the smallest. Each inner circle represents the genome sequence of an isolate (1 to 13). Each radial line indicates the position of an SNV on the scaffold. The color of each radial line corresponds to the genotype of the SNV, with the following color code: green = heterozygous state (genotype "0/1"); black = homozygous state with the reference haplotype (genotype "0/0"); red = homozygous state with the alternative haplotype (genotype "1/1"). Therefore, “green” regions indicate heterozygosity, while “red” and “black” regions indicate loss of heterozygosity (LoH). The blue regions represent conserved homozygous regions.

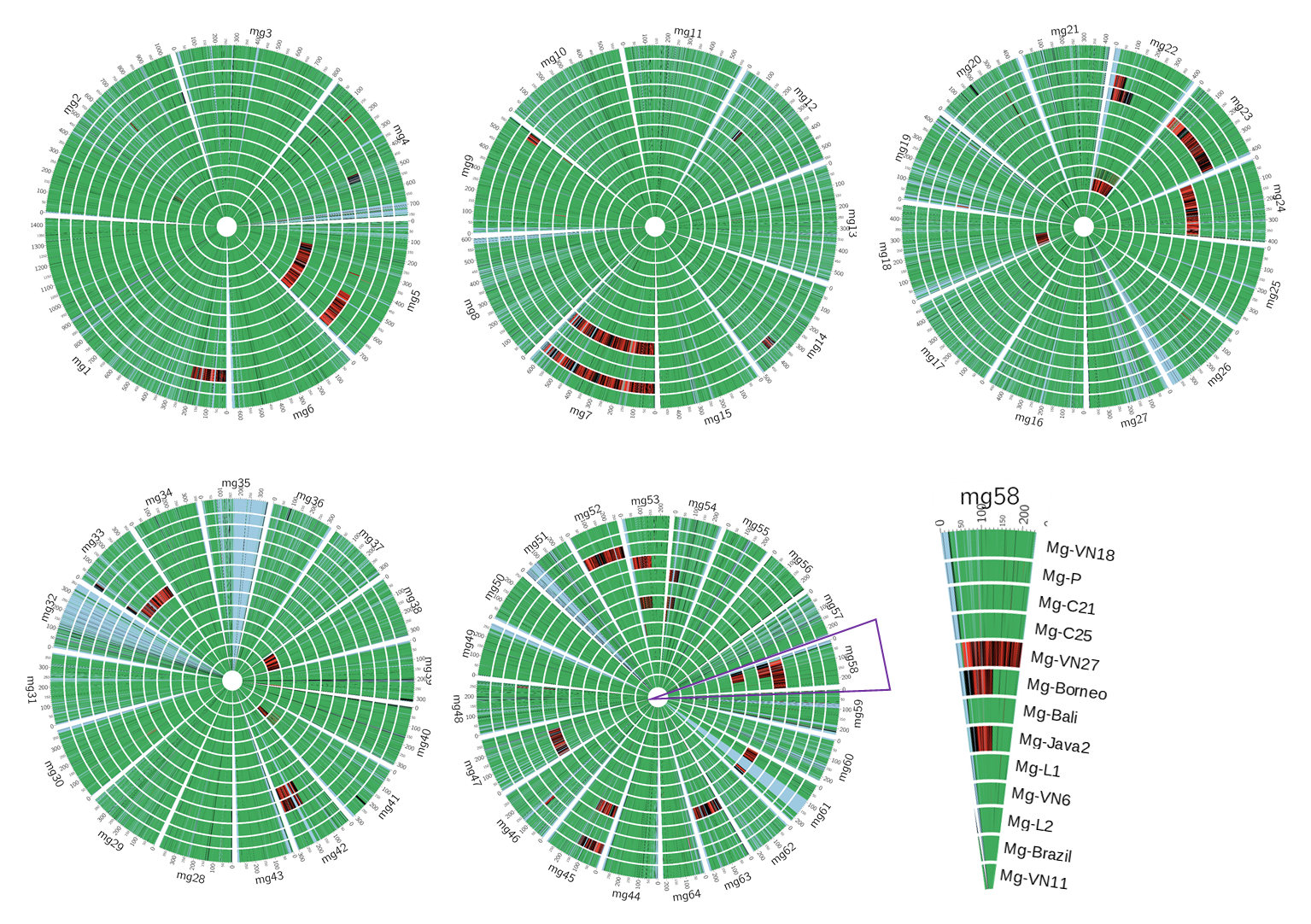
