## Supplementary Figure S3 for "Genomic rearrangements promote diversification of a facultative meiotic parthenogenetic nematode pest (*Meloidogyne graminicola*)"

Heterozygosity levels of the genomes of 13 isolates based on *k*-mer analysis

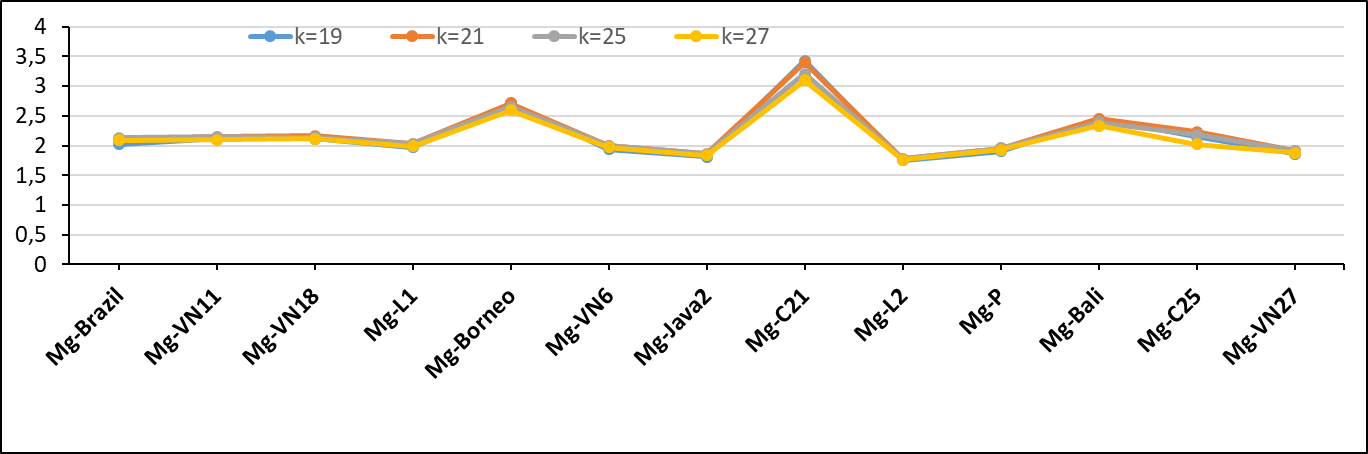
