## Supplementary Figure S4 for "Genomic rearrangements promote diversification of a facultative meiotic parthenogenetic nematode pest (*Meloidogyne graminicola*)"

### Linkage disequilibrium and 4‐gametes test of *M. graminicola* isolates using the 40 fixed markers. The red line represents the r^2^ correlation between markers, indicating linkage disequilibrium (LD) for each physical distance between the SNV markers. The blue line represents the proportion of pairs of two‐state markers that pass the 4‐gamete test for each physical distance between the markers on a scaffold.

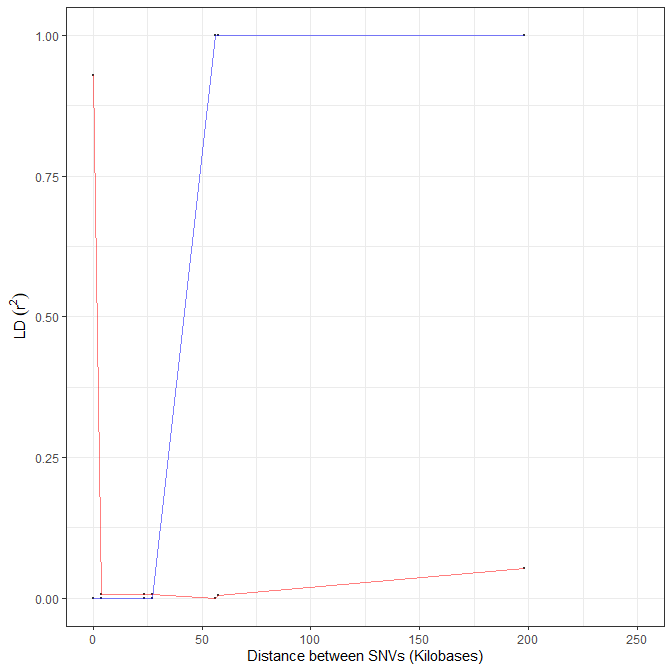
