## Supplementary Figure S5 for "Genomic rearrangements promote diversification of a facultative meiotic parthenogenetic nematode pest (*Meloidogyne graminicola*)"

### Haplotype networks (A) and principal component analysis (B) among *M. graminicola* isolates using the fixed 40 SNVs. A) Code names of isolates are indicated in the circles and countries of origin are displayed by different colors. Each slash on the branches represents a SNV.

**(A)**

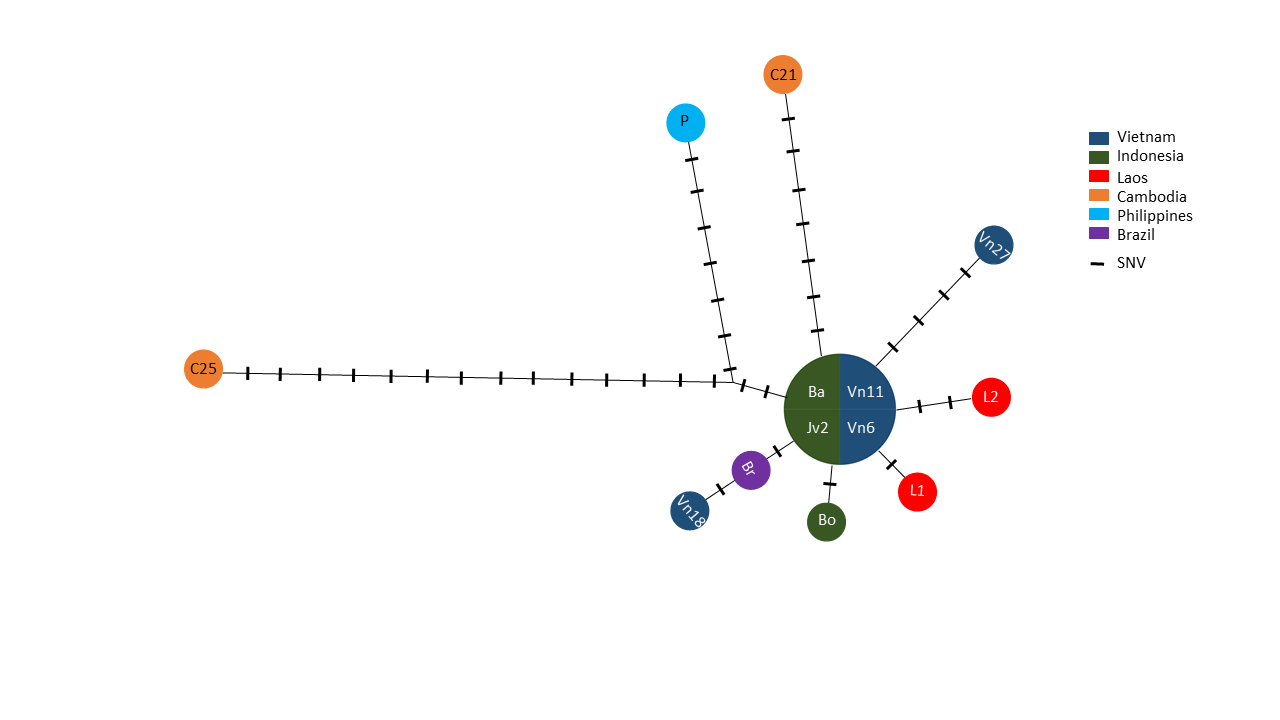

**(B)**

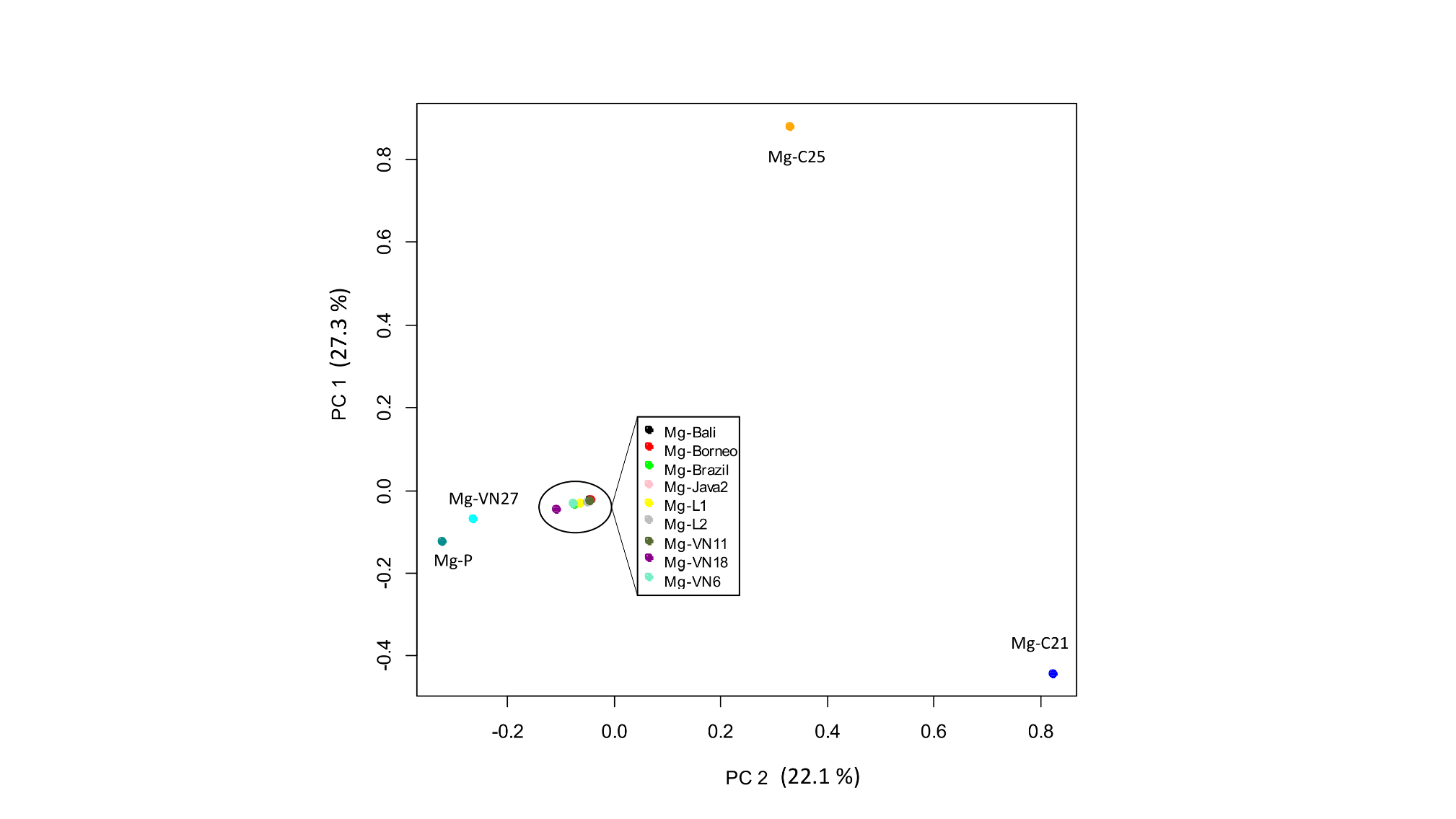
