## Supplementary Figure S6 for "Genomic rearrangements promote diversification of a facultative meiotic parthenogenetic nematode pest (*Meloidogyne graminicola*)"

### Per-base coverage analysis of HiSeq data from the Mg-VN18 isolate on the consensus sequence of the GB2 contig and two divergent copies (GB2_T1 and GB2_T2). Grey cloud indicates coverage of the reads, which are identical to the sequence template (GB2_T1, GB2_T2). The phased colors in the first track correspond to coverage of the different reads (belong to two divergent haplotypes) at the same position of consensus sequence (GB2). The number on the left side of each track indicates average coverage of reads on each fragment.

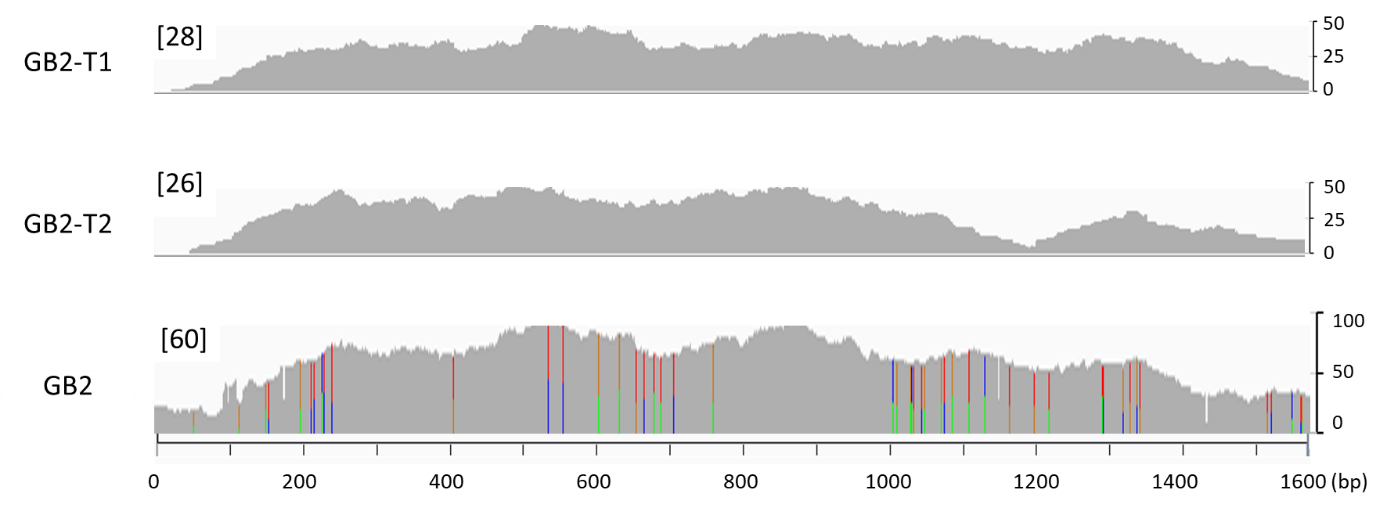
