## Supplementary Figure S7 for "Genomic rearrangements promote diversification of a facultative meiotic parthenogenetic nematode pest (*Meloidogyne graminicola*)"

### Average read-coverage on the two haplotypes of 18 contigs (on the x axis) among the 13 *M. graminicola* isolates (on the right y axis). The black horizontal line indicates mean value of read-coverage of all divergent copies in each isolate. Missing columns (read-coverage = 0×) indicates the absence of one haplotype.

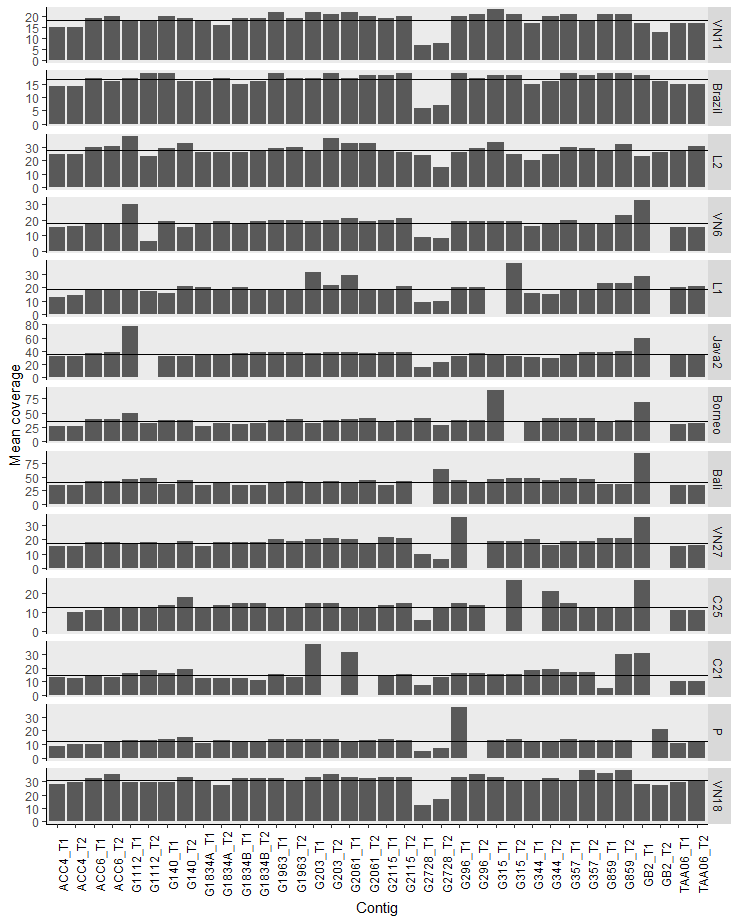
