## Supplementary Figure S8 for "Genomic rearrangements promote diversification of a facultative meiotic parthenogenetic nematode pest (*Meloidogyne graminicola*)"

### Average read-coverage on 36 divergent copies of 18 contigs using full reads (read-depth ranged from 288 to 568×) in five isolates (Mg-VN18, Mg-L2, Mg-Borneo, Mg-Bali, Mg-Java2). The black horizontal line indicates the mean value of read-coverage of all divergent copies in each isolate. The missing columns indicate the absence of copies (average read-coverage = 0×).

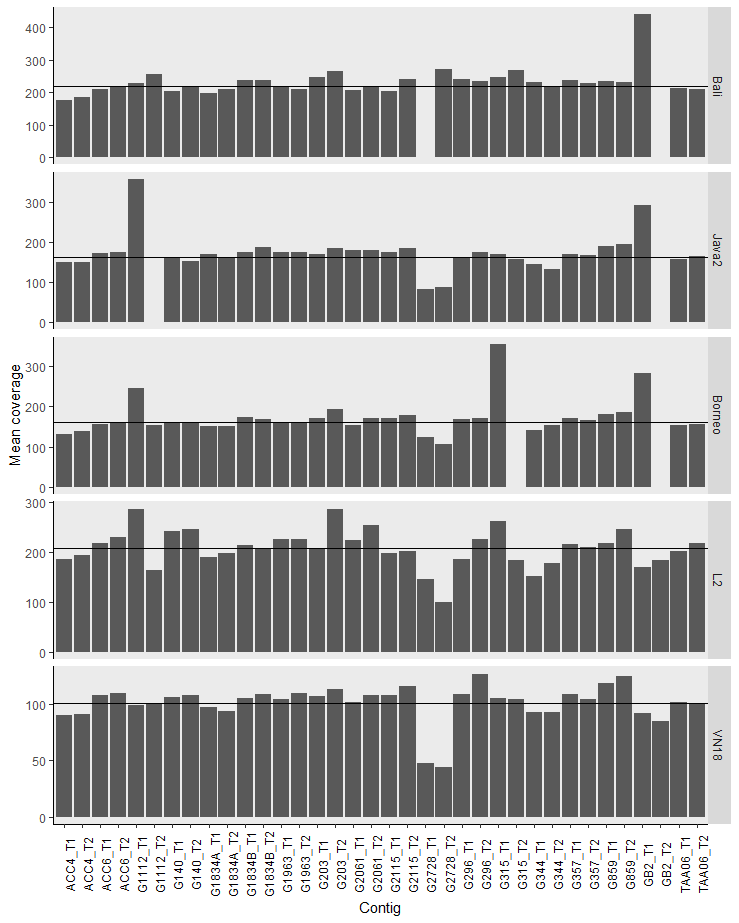
