## Supplementary Figure S9 for "Genomic rearrangements promote diversification of a facultative meiotic parthenogenetic nematode pest (*Meloidogyne graminicola*)"

Distribution of loss of heterozygosity events (LoH), duplication (DUP), and deletion (DEL) identified in 142 scaffolds among 13 *Mg* isolates. The lengths of the scaffolds are presented to scale. Each horizontal line represents the LoH events of an isolate. Each eclipse and triangle indicate the duplication event and deletion of an isolate, respectively. The colors correspond to isolates

| 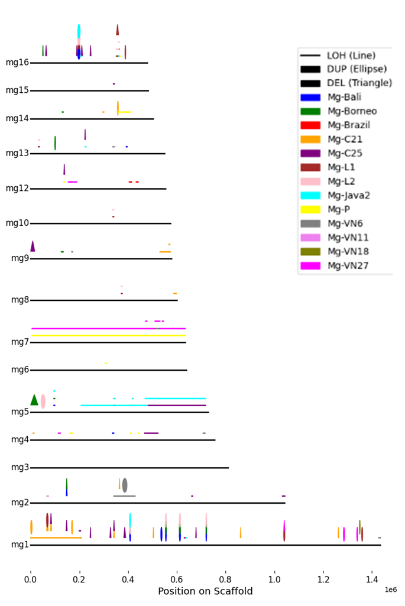 | 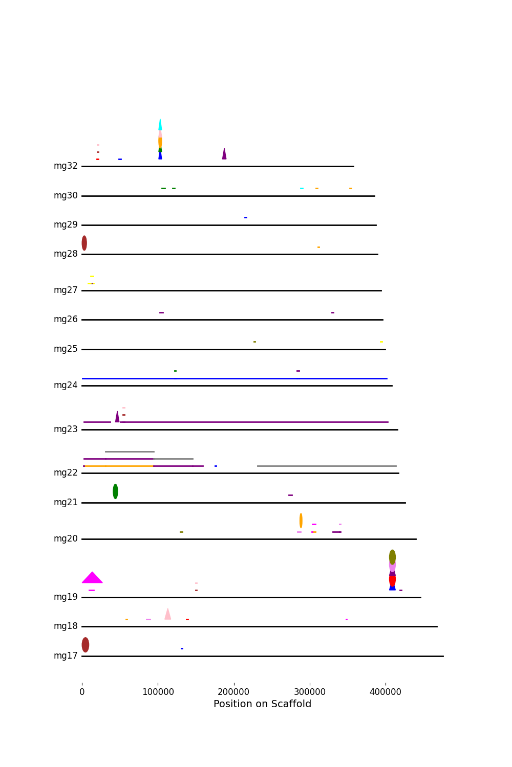 | 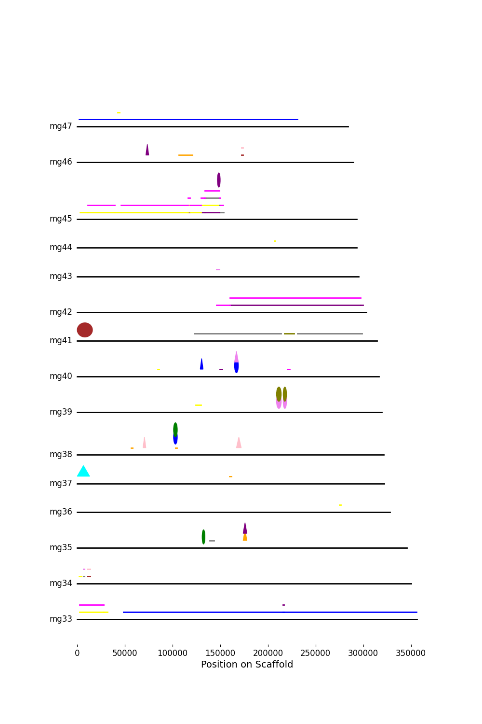 | 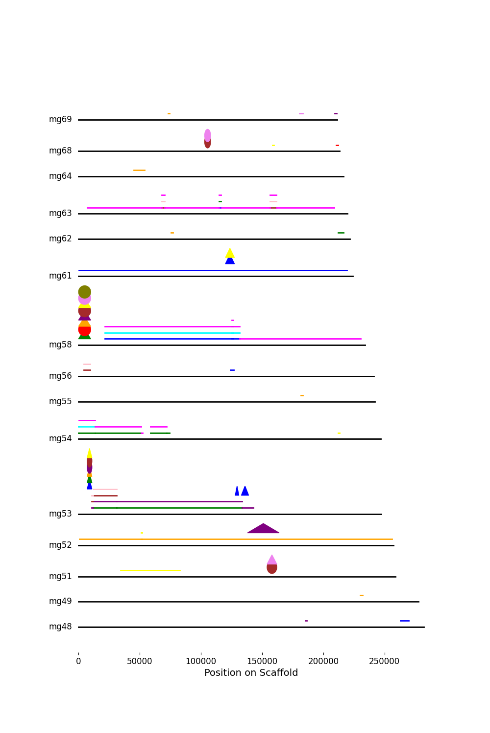 | 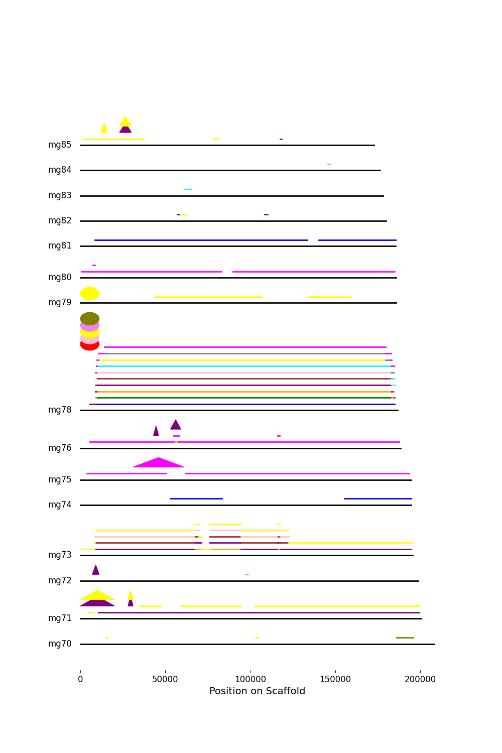 |
| --- | --- | --- | --- | --- |
| 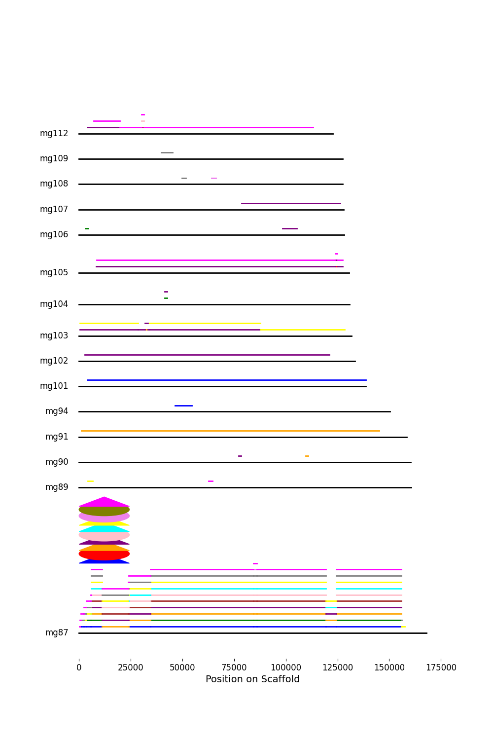 | 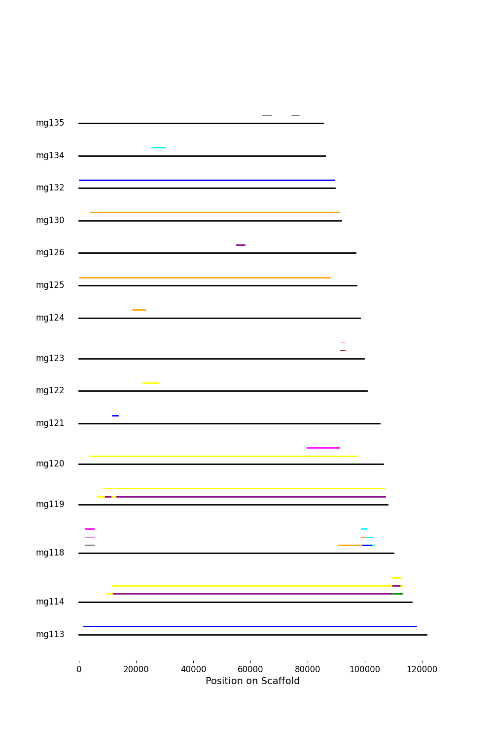 | 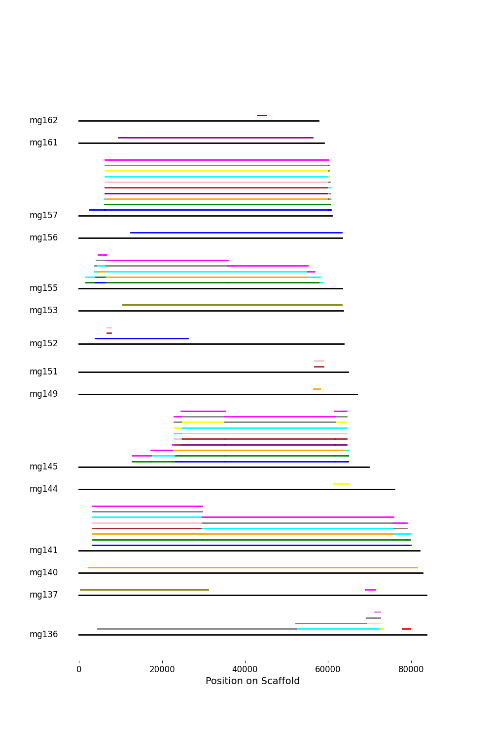 | 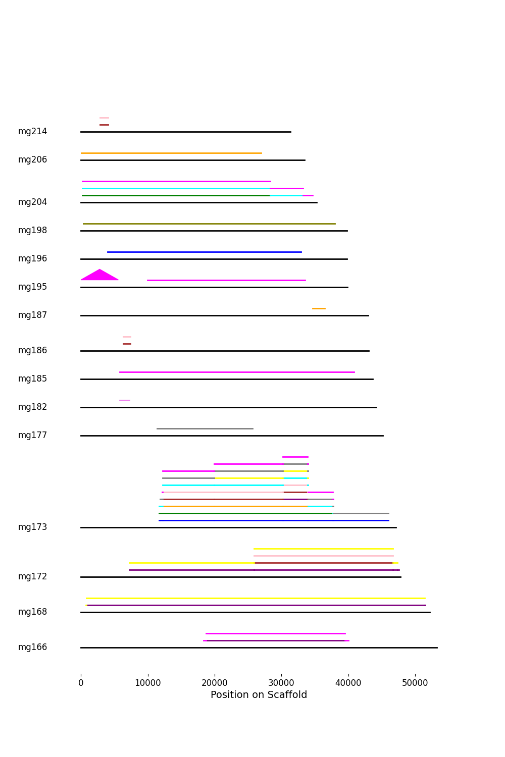 | 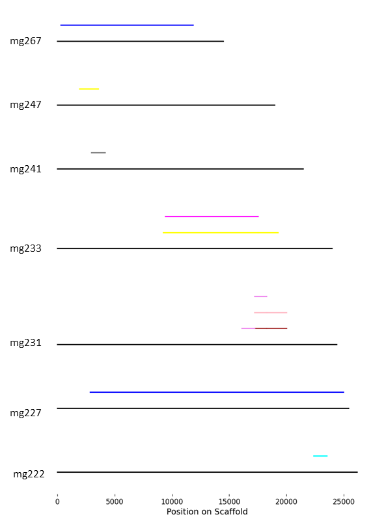 |
