## Supplementary Table S1 for "Genomic rearrangements promote diversification of a facultative meiotic parthenogenetic nematode pest (*Meloidogyne graminicola*)"

### Description of the *Meloidogyne graminicola* isolates used in this study and of the generated genomic data: number of raw and cleaned paired end Illumina reads (# raw PE reads and # cleaned PE reads, respectively), read length, GC content of the assembly (%GC), and mean coverage (Coverage).

| No | Population | Country | # raw PE reads | # cleaned PE reads | Read length | %GC | Coverage**^2^** |
| --- | --- | --- | --- | --- | --- | --- | --- |
| 1 | Mg-VN18**^1^** | Vietnam | 122,890,657 | 86,991,142 | 150 | 24 | 288 |
| 2 | Mg-Borneo | Indonesia | 147,469,360 | 99,625,819 | 150 | 24 | 360 |
| 3 | Mg-Bali | Indonesia | 180,145,846 | 122,418,010 | 150 | 24 | 442 |
| 4 | Mg-Java2 | Indonesia | 118,708,848 | 118,590,640 | 150 | 24 | 420 |
| 5 | Mg-L2 | Laos | 157,229,694 | 156,871,340 | 150 | 24 | 568 |
| 6 | Mg-L1 | Laos | 13,807,048 | 15,746,640 | 125 | 26 | 38 |
| 7 | Mg-P | Philippines | 15,524,276 | 9,867,172 | 100 | 24 | 24 |
| 8 | Mg-Brazil | Brazil | 16,880,430 | 16,339,898 | 100 | 26 | 39 |
| 9 | Mg-C21 | Cambodia | 17,457,706 | 8,637,052 | 125 | 24 | 26 |
| 10 | Mg-C25 | Cambodia | 26,347,326 | 7,045,100 | 125 | 25 | 21 |
| 11 | Mg-VN6 | Vietnam | 16,374,636 | 15,182,532 | 100 | 24 | 37 |
| 12 | Mg-VN11 | Vietnam | 17,227,778 | 15,746,640 | 100 | 25 | 38 |
| 13 | Mg-VN27 | Vietnam | 16,298,908 | 15,137,918 | 100 | 24 | 36 |

^1^ The genome of isolate Mg-VN18 was used to generate the haploid reference genome sequence in Phan et al. [5]. ^2^ Average read coverage considering assembly genome size of 41.5 Mb.

REFERENCES

5. Phan NT, Orjuela J, Danchin EGJ, Klopp C, Perfus-Barbeoch L, Kozlowski DK et al. Genome structure and content of the rice root-knot nematode (*Meloidogyne graminicola*). Ecol Evol.
