## Supplementary Table S2 for "Genomic rearrangements promote diversification of a facultative meiotic parthenogenetic nematode pest (*Meloidogyne graminicola*)"

### Description of the 18 manually curated contigs (covering 177 kb) with a comparison between their two sequence types (haplotypes)

| No | Contig name | Type | Divergent copy name | Length (bp) | Position on reference genome**^1^** | # Genes | Identity between 2 types |
| --- | --- | --- | --- | --- | --- | --- | --- |
| 1 | ACC4 | Type 1 | ACC4_T1 | 11,247 | mg4:505760-516936 | 2 | 94.21 |
|  |  | Type 2 | ACC4_T2 | 11,214 |  |  |  |
| 2 | G140 | Type 1 | G140_T1 | 1,429 | mg6:134432-135865 | 1 | 96.66 |
|  |  | Type 2 | G140_T2 | 1,433 |  |  |  |
| 3 | G296 | Type 1 | G296_T1 | 12,282 | mg7:6227-18522 | 2 | 97.67 |
|  |  | Type 2 | G296_T2 | 12,323 |  |  |  |
| 4 | ACC6**^2^** | Type 1 | ACC6_T1 | 6,310 | mg15:201656-207657 | 1 | 94.73 |
|  |  | Type 2 | ACC6_T2 | 6,151 |  |  |  |
| 5 | G1963 | Type 1 | G1963_T1 | 8,687 | mg25:79960-88655 | 5 | 96.56 |
|  |  | Type 2 | G1963_T2 | 8,690 |  |  |  |
| 6 | G1834A | Type 1 | G1834A_T1 | 2,240 | mg44:37041-39278 | 1 | 99.11 |
|  |  | Type 2 | G1834A_T2 | 2,238 |  |  |  |
| 7 | G1834B | Type 1 | G1834B_T1 | 3,110 | mg44:39200-42308 | 1 | 99.71 |
|  |  | Type 2 | G1834B_T2 | 3,111 |  |  |  |
| 8 | G315 | Type 1 | G315_T1 | 6,258 | mg53:17740-24025 | 1 | 97.73 |
|  |  | Type 2 | G315_T2 | 6,282 |  |  |  |
| 9 | TAA06**^2^** | Type 1 | TAA06_T1 | 6,361 | mg68:90615-96976 | 3 | 96.05 |
|  |  | Type 2 | TAA06_T2 | 6,399 |  |  |  |
| 10 | GB2 | Type 1 | GB2_T1 | 1,589 | mg78:21353-22923 | 1 | 94.99 |
|  |  | Type 2 | GB2_T2 | 1,572 |  |  |  |
| 11 | G2115 | Type 1 | G2115_T1 | 7,868 | mg93:111858-119733 | 1 | 98.19 |
|  |  | Type 2 | G2115_T2 | 7,865 |  |  |  |
| 12 | G2061 | Type 1 | G2061_T1 | 2,897 | mg125:29921-32815 | 1 | 95.22 |
|  |  | Type 2 | G2061_T2 | 2,906 |  |  |  |
| 13 | G203 | Type 1 | G203_T1 | 17,866 | mg125:68032-85564 | 3 | 98.50 |
|  |  | Type 2 | G203_T2 | 17,172 |  |  |  |
| 14 | G859 | Type 1 | G859_T1 | 80,275 | mg130:4438-84558 | 26 | 97.42 |
|  |  | Type 2 | G859_T2 | 79,436 |  |  |  |
| 15 | G1112 | Type 1 | G1112_T1 | 2,810 | mg136:62364-65160 | 1 | 97.48 |
|  |  | Type 2 | G1112_T2 | 2,800 |  |  |  |
| 16 | G2728 | Type 1 | G2728_T1 | 532 | mg156:12687-13219 | 2 | 98.08 |
|  |  | Type 2 | G2728_T2 | 526 |  |  |  |
| 17 | G344 | Type 1 | G344_T1 | 2,065 | mg161:11242-13302 | 1 | 97.83 |
|  |  | Type 2 | G344_T2 | 2,061 |  |  |  |
| 18 | G357 | Type 1 | G357_T1 | 3,904 | mg199:8629-12519 | 1 | 96.64 |
|  |  | Type 2 | G357_T2 | 3,888 |  |  |  |

^1^scaffold name: start position-end position

^2^two contigs from Besnard et al. [2]

REFERENCES

2. Besnard G, Thi-Phan N, Ho-Bich H, Dereeper A, Nguyen HT, Quénéhervé P, et al. On the close relatedness of two rice-parasitic root-knot nematode species and the recent expansion of *Meloidogyne graminicola* in Southeast Asia. Genes (Basel). 2019;10(2). doi: 10.3390/genes10020175
