## Supplementary Table S3 for "Genomic rearrangements promote diversification of a facultative meiotic parthenogenetic nematode pest (*Meloidogyne graminicola*)"

### Number of regions bearing copy number variants (CNV), number of genes overlap with CNV regions (at least 70% length), and number of putative secreted protein coding genes located on CNV regions in 13 *M. graminicola* isolates. CN0 and CN1 indicate complete deletion and single deletion of a copy, respectively, compared to normal diploid state; CN3 to CN6 indicate the number of copies (from three to six copies) compared to the normal diploid state.

| #copy | Number of CNV regions | | | | | | | | | | | | |
| --- | --- | --- | --- | --- | --- | --- | --- | --- | --- | --- | --- | --- | --- |
|  | Mg-Bali | Mg-Borneo | Mg-Brazil | Mg-C21 | Mg-C25 | Mg-L1 | Mg-L2 | Mg-Java2 | Mg-P | Mg-VN6 | Mg-VN11 | Mg-VN18 | Mg-VN27 |
| Deletion (CN0) | 14 | 2 | 0 | 9 | 24 | 0 | 5 | 0 | 21 | 0 | 0 | 0 | 36 |
| Half deletion (CN1) | 20 | 30 | 0 | 22 | 75 | 11 | 4 | 11 | 13 | 0 | 7 | 0 | 1 |
| Three copies (CN3) | 12 | 8 | 10 | 11 | 2 | 38 | 17 | 5 | 0 | 12 | 16 | 13 | 2 |
| Four copies (CN4) | 0 | 1 | 0 | 2 | 2 | 9 | 0 | 0 | 4 | 0 | 0 | 0 | 0 |
| Five copies (CN5) | 2 | 0 | 0 | 0 | 0 | 0 | 0 | 0 | 0 | 0 | 0 | 0 | 0 |
| Six copies (CN6) | 0 | 0 | 0 | 0 | 0 | 2 | 2 | 0 | 2 | 0 | 2 | 2 | 0 |
| #copy | Number of genes which have length overlap at least 70% with CNV regions | | | | | | | | | | | | |
| Deletion (CN0) | 8 | 2 | 0 | 7 | 20 | 0 | 4 | 0 | 17 | 0 | 0 | 0 | 24 |
| Half deletion (CN1) | 18 | 18 | 0 | 21 | 59 | 11 | 3 | 9 | 9 | 0 | 7 | 0 | 1 |
| Three copies (CN3) | 10 | 6 | 8 | 11 | 2 | 34 | 15 | 5 | 0 | 10 | 14 | 11 | 2 |
| Four copies (CN4) | 0 | 1 | 0 | 2 | 2 | 9 | 0 | 0 | 4 | 0 | 0 | 0 | 0 |
| Five copies (CN5) | 2 | 0 | 0 | 0 | 0 | 0 | 0 | 0 | 0 | 0 | 0 | 0 | 0 |
| Six copies (CN6) | 0 | 0 | 0 | 0 | 0 | 2 | 2 | 0 | 2 | 0 | 2 | 2 | 0 |
| #copy | Number of putative secreted protein coding genes located on CNV regions | | | | | | | | | | | | |
| Deletion (CN0) | 0 | 0 | 0 | 0 | 2 | 0 | 0 | 0 | 0 | 0 | 0 | 0 | 8 |
| Half deletion (CN1) | 0 | 0 | 0 | 0 | 10 | 0 | 0 | 0 | 1 | 0 | 0 | 0 | 0 |
| Three copies (CN3) | 2 | 3 | 0 | 0 | 0 | 2 | 1 | 0 | 0 | 0 | 0 | 0 | 0 |
| Four copies (CN4) | 0 | 0 | 0 | 0 | 0 | 0 | 0 | 0 | 1 | 0 | 0 | 0 | 0 |
