## Supplementary Table S4 for "Genomic rearrangements promote diversification of a facultative meiotic parthenogenetic nematode pest (*Meloidogyne graminicola*)"

### Name and domain functions of 27 putative secreted protein-coding genes located in CNV regions among 13 isolates

| #Copy | Gene ID | Domains | | General process/function |
| --- | --- | --- | --- | --- |
| CN0 | mg19g0046821 |  | Unknown |  |
|  | mg19g0046831 mg19g0046871 | IPR004898 | Pectate lyase PlyH/PlyE-like Pectin lyase fold/virulence factor | Plant cell-wall degradation |
|  |  | IPR012334 |  |  |
|  |  | IPR011050 |  |  |
|  | mg19g0046881 |  | Unknown |  |
|  | mg23g0053771 |  | Unknown |  |
|  | mg75g0107381 |  | Unknown |  |
|  | mg75g0107401 |  | Unknown |  |
|  | mg75g0107421 | IPR033112 | Phospholipase A2, aspartic acid active site | Modulation of host defense mechanisms |
|  |  | IPR036444 |  |  |
|  |  | IPR016090 |  |  |
|  | mg75g0107451 | IPR038577 | C-terminal domain superfamily | Modification of nematode effectors |
|  |  | IPR001503 | Glycosyl transferase family 10 |  |
|  |  | IPR031481 | Fucosyltransferase, N-terminal |  |
|  | mg76g0108031 | IPR006026 | Peptidase, metallopeptidase | Protein degradation |
|  |  | IPR001506 | Peptidase M12A |  |
|  |  | IPR024079 | Metallopeptidase |  |
|  |  | IPR034035 | Astacin-like metallopeptidase domain |  |
| CN1 | mg1g0001561 | IPR024079 | Metallopeptidase |  |
|  | mg9g0025021 | IPR035595 | UDP-glucuronosyl | Modulation of hot defense mechanism |
|  |  | IPR002213 | UDP-glucosyltransferase |  |
|  | mg9g0025041 |  | Unknown |  |
|  | mg31g0066071 |  | Unknown |  |
|  | mg31g0066081 | IPR011050 | Pectin lyase fold/virulence factor RCR-like domain superfamily Parallel beta-heli Unknown repeat | Plant cell-wall degradation |
|  |  | IPR036772 |  |  |
|  |  | IPR012334 |  |  |
|  |  | IPR001190 |  |  |
|  |  | IPR017448 |  |  |
|  |  | IPR006626 |  |  |
|  | mg31g0066131 |  | Unknown |  |
|  | mg52g0090041 |  | Unknown |  |
|  | mg52g0090051 | IPR001828 | Receptor ligand binding region | Host recognition and attachment |
|  |  | IPR028082 | Periplasmic binding protein-like I |  |
|  | mg85g0113571 |  | Unknown |  |
|  | mg92g0117091 |  | Unknown |  |
|  | mg92g0117111 |  | Unknown |  |
| CN3 | mg1g0002381 | IPR001507 | Zona pellucida domain | Modulation of host defense mechanisms |
|  | mg1g0005091 |  | Unknown |  |
|  | mg21g0050091 | IPR036084 | Serine protease inhibitor-like | Inhibiting host plant proteases |
|  |  | IPR002919 | Trypsin Inhibitor-like,  cysteine rich domain |  |
|  | mg38g0073821 |  | Unknown |  |
|  | mg77g0108621 | IPR019330 | LRP chaperone MESD | Interactions with host plant cells/ Feeding site formation |
| CN4 | mg79g0110191 | IPR000742 | EGF-like domain |  |
