## Supplementary Table S5 for "Genomic rearrangements promote diversification of a facultative meiotic parthenogenetic nematode pest (*Meloidogyne graminicola*)"

### Details of CNVs found on scaffolds among 13 isolates. CN2 indicates normal diploid state; CN0 and CN1 indicate complete deletion or single deletion compared to the normal diploid state; CN3 to CN6 indicate higher copy number compared to the normal diploid state.

|  | Scaffolds | Start | End | Width | Number of copies | | | | | | | | | | | | |
| --- | --- | --- | --- | --- | --- | --- | --- | --- | --- | --- | --- | --- | --- | --- | --- | --- | --- |
|  |  |  |  |  | Mg-Bali | Mg-Borneo | Mg-Brazil | Mg-C21 | Mg-C25 | Mg-L1 | Mg-L2 | Mg-Java2 | Mg-P | Mg-VN6 | Mg-VN11 | Mg-VN18 | Mg-VN27 |
| 1 | mg1 | 0 | 6000 | 6001 | CN2 | CN2 | CN2 | CN3 | CN2 | CN2 | CN2 | CN2 | CN2 | CN2 | CN2 | CN2 | CN2 |
| 2 | mg1 | 65000 | 72000 | 7001 | CN2 | CN2 | CN2 | CN0 | CN2 | CN4 | CN2 | CN2 | CN2 | CN2 | CN2 | CN2 | CN2 |
| 3 | mg1 | 81000 | 86000 | 5001 | CN2 | CN2 | CN2 | CN1 | CN1 | CN2 | CN2 | CN2 | CN2 | CN2 | CN2 | CN2 | CN2 |
| 4 | mg1 | 145000 | 150000 | 5001 | CN2 | CN2 | CN2 | CN2 | CN1 | CN2 | CN2 | CN2 | CN2 | CN2 | CN2 | CN2 | CN2 |
| 5 | mg1 | 168000 | 172000 | 4001 | CN2 | CN2 | CN2 | CN4 | CN2 | CN2 | CN2 | CN2 | CN2 | CN2 | CN2 | CN2 | CN2 |
| 6 | mg1 | 244000 | 247000 | 3001 | CN2 | CN2 | CN2 | CN2 | CN1 | CN2 | CN2 | CN2 | CN2 | CN2 | CN2 | CN2 | CN2 |
| 7 | mg1 | 325000 | 330000 | 5001 | CN2 | CN2 | CN2 | CN2 | CN1 | CN2 | CN2 | CN2 | CN2 | CN2 | CN2 | CN2 | CN2 |
| 8 | mg1 | 340000 | 344000 | 4001 | CN2 | CN2 | CN2 | CN1 | CN1 | CN2 | CN2 | CN2 | CN2 | CN2 | CN2 | CN2 | CN2 |
| 9 | mg1 | 382000 | 390000 | 8001 | CN2 | CN2 | CN2 | CN2 | CN1 | CN2 | CN2 | CN2 | CN2 | CN2 | CN2 | CN2 | CN2 |
| 10 | mg1 | 406000 | 410000 | 4001 | CN3 | CN2 | CN2 | CN2 | CN2 | CN2 | CN3 | CN3 | CN2 | CN2 | CN2 | CN2 | CN2 |
| 11 | mg1 | 502000 | 505000 | 3001 | CN2 | CN2 | CN2 | CN1 | CN2 | CN2 | CN2 | CN2 | CN2 | CN2 | CN2 | CN2 | CN2 |
| 12 | mg1 | 534000 | 539000 | 5001 | CN3 | CN2 | CN2 | CN2 | CN2 | CN2 | CN2 | CN2 | CN2 | CN2 | CN2 | CN2 | CN2 |
| 13 | mg1 | 553000 | 557000 | 4001 | CN3 | CN3 | CN2 | CN2 | CN2 | CN2 | CN3 | CN2 | CN2 | CN2 | CN2 | CN2 | CN2 |
| 14 | mg1 | 610000 | 615000 | 5001 | CN3 | CN3 | CN2 | CN2 | CN2 | CN2 | CN3 | CN2 | CN2 | CN2 | CN2 | CN2 | CN2 |
| 15 | mg1 | 677000 | 680000 | 3001 | CN2 | CN2 | CN2 | CN2 | CN1 | CN2 | CN2 | CN2 | CN2 | CN2 | CN2 | CN2 | CN2 |
| 16 | mg1 | 719000 | 722000 | 3001 | CN3 | CN3 | CN2 | CN2 | CN2 | CN2 | CN3 | CN2 | CN2 | CN2 | CN2 | CN2 | CN2 |
| 17 | mg1 | 860000 | 863000 | 3001 | CN2 | CN2 | CN2 | CN1 | CN2 | CN2 | CN2 | CN2 | CN2 | CN2 | CN2 | CN2 | CN2 |
| 18 | mg1 | 1038000 | 1043000 | 5001 | CN2 | CN2 | CN2 | CN2 | CN2 | CN3 | CN2 | CN2 | CN2 | CN2 | CN2 | CN2 | CN1 |
| 19 | mg1 | 1261000 | 1264000 | 3001 | CN2 | CN2 | CN2 | CN1 | CN2 | CN2 | CN2 | CN2 | CN2 | CN2 | CN2 | CN2 | CN2 |
| 20 | mg1 | 1283000 | 1288000 | 5001 | CN2 | CN2 | CN2 | CN2 | CN2 | CN2 | CN2 | CN2 | CN2 | CN2 | CN2 | CN2 | CN3 |
| 21 | mg1 | 1338000 | 1342000 | 4001 | CN2 | CN2 | CN2 | CN2 | CN2 | CN2 | CN2 | CN2 | CN2 | CN2 | CN2 | CN2 | CN3 |
| 22 | mg1 | 1349000 | 1352000 | 3001 | CN2 | CN2 | CN2 | CN2 | CN2 | CN2 | CN2 | CN2 | CN2 | CN2 | CN2 | CN3 | CN2 |
| 23 | mg1 | 1358000 | 1362000 | 4001 | CN2 | CN2 | CN2 | CN2 | CN2 | CN3 | CN2 | CN2 | CN2 | CN2 | CN2 | CN2 | CN2 |
| 24 | mg2 | 363000 | 366000 | 3001 | CN2 | CN2 | CN2 | CN1 | CN2 | CN2 | CN2 | CN2 | CN2 | CN2 | CN2 | CN2 | CN2 |
| 25 | mg2 | 377000 | 395552 | 18553 | CN2 | CN2 | CN2 | CN2 | CN2 | CN2 | CN2 | CN2 | CN2 | CN3 | CN2 | CN2 | CN2 |
| 26 | mg2 | 146000 | 150000 | 4001 | CN1 | CN1 | CN2 | CN2 | CN2 | CN2 | CN2 | CN2 | CN2 | CN2 | CN2 | CN2 | CN2 |
| 27 | mg5 | 0 | 30000 | 30001 | CN2 | CN1 | CN2 | CN2 | CN2 | CN2 | CN2 | CN2 | CN2 | CN2 | CN2 | CN2 | CN2 |
| 28 | mg5 | 43000 | 59000 | 16001 | CN2 | CN2 | CN2 | CN2 | CN2 | CN2 | CN3 | CN2 | CN2 | CN2 | CN2 | CN2 | CN2 |
| 29 | mg9 | 0 | 16893 | 16894 | CN2 | CN2 | CN2 | CN2 | CN1 | CN2 | CN2 | CN2 | CN2 | CN2 | CN2 | CN2 | CN2 |
| 30 | mg12 | 136000 | 140000 | 4001 | CN2 | CN2 | CN2 | CN2 | CN1 | CN2 | CN2 | CN2 | CN2 | CN2 | CN2 | CN2 | CN2 |
| 31 | mg13 | 99000 | 102000 | 3001 | CN2 | CN4 | CN2 | CN2 | CN2 | CN2 | CN2 | CN2 | CN2 | CN2 | CN2 | CN2 | CN2 |
| 32 | mg13 | 222000 | 225000 | 3001 | CN2 | CN2 | CN2 | CN2 | CN1 | CN2 | CN2 | CN2 | CN2 | CN2 | CN2 | CN2 | CN2 |
| 33 | mg14 | 356000 | 360000 | 4001 | CN2 | CN2 | CN2 | CN3 | CN2 | CN2 | CN2 | CN2 | CN2 | CN2 | CN2 | CN2 | CN2 |
| 34 | mg16 | 49000 | 52000 | 3001 | CN2 | CN1 | CN2 | CN2 | CN2 | CN2 | CN2 | CN2 | CN2 | CN2 | CN2 | CN2 | CN2 |
| 35 | mg16 | 61000 | 66000 | 5001 | CN2 | CN2 | CN2 | CN2 | CN1 | CN2 | CN2 | CN2 | CN2 | CN2 | CN2 | CN2 | CN2 |
| 36 | mg16 | 188000 | 191000 | 3001 | CN2 | CN2 | CN2 | CN2 | CN1 | CN2 | CN2 | CN2 | CN2 | CN2 | CN2 | CN2 | CN2 |
| 37 | mg16 | 193000 | 203000 | 10001 | CN3 | CN2 | CN2 | CN2 | CN2 | CN1 | CN3 | CN3 | CN2 | CN2 | CN2 | CN2 | CN2 |
| 38 | mg16 | 207000 | 213000 | 6001 | CN2 | CN2 | CN2 | CN2 | CN1 | CN2 | CN2 | CN2 | CN2 | CN2 | CN2 | CN2 | CN2 |
| 39 | mg16 | 244000 | 248000 | 4001 | CN2 | CN2 | CN2 | CN2 | CN1 | CN2 | CN2 | CN2 | CN2 | CN2 | CN2 | CN2 | CN2 |
| 40 | mg16 | 352000 | 361000 | 9001 | CN2 | CN2 | CN2 | CN2 | CN2 | CN1 | CN2 | CN2 | CN2 | CN2 | CN2 | CN2 | CN2 |
| 41 | mg16 | 387000 | 390000 | 3001 | CN2 | CN2 | CN2 | CN2 | CN2 | CN3 | CN2 | CN2 | CN2 | CN2 | CN2 | CN2 | CN2 |
| 42 | mg17 | 0 | 9000 | 9001 | CN2 | CN2 | CN2 | CN2 | CN2 | CN4 | CN2 | CN2 | CN2 | CN2 | CN2 | CN2 | CN2 |
| 43 | mg18 | 109000 | 117000 | 8001 | CN2 | CN2 | CN2 | CN2 | CN2 | CN2 | CN0 | CN2 | CN2 | CN2 | CN2 | CN2 | CN2 |
| 44 | mg19 | 405000 | 413000 | 8001 | CN1 | CN2 | CN3 | CN2 | CN0 | CN2 | CN2 | CN2 | CN2 | CN2 | CN3 | CN3 | CN2 |
| 45 | mg19 | 0 | 27000 | 27001 | CN2 | CN2 | CN2 | CN2 | CN2 | CN2 | CN2 | CN2 | CN2 | CN2 | CN2 | CN2 | CN0 |
| 46 | mg20 | 287000 | 290000 | 3001 | CN2 | CN2 | CN2 | CN3 | CN2 | CN2 | CN2 | CN2 | CN2 | CN2 | CN2 | CN2 | CN2 |
| 47 | mg21 | 41000 | 47000 | 6001 | CN2 | CN3 | CN2 | CN2 | CN2 | CN2 | CN2 | CN2 | CN2 | CN2 | CN2 | CN2 | CN2 |
| 48 | mg23 | 44000 | 49000 | 5001 | CN2 | CN2 | CN2 | CN2 | CN0 | CN2 | CN2 | CN2 | CN2 | CN2 | CN2 | CN2 | CN2 |
| 49 | mg28 | 0 | 6000 | 6001 | CN2 | CN2 | CN2 | CN2 | CN2 | CN3 | CN2 | CN2 | CN2 | CN2 | CN2 | CN2 | CN2 |
| 50 | mg31 | 37000 | 44000 | 7001 | CN2 | CN2 | CN2 | CN2 | CN1 | CN2 | CN2 | CN2 | CN2 | CN2 | CN2 | CN2 | CN2 |
| 51 | mg31 | 55000 | 59000 | 4001 | CN2 | CN2 | CN2 | CN2 | CN1 | CN2 | CN2 | CN2 | CN2 | CN2 | CN2 | CN2 | CN2 |
| 52 | mg32 | 101000 | 105000 | 4001 | CN1 | CN1 | CN2 | CN4 | CN2 | CN2 | CN1 | CN1 | CN2 | CN2 | CN2 | CN2 | CN2 |
| 53 | mg32 | 185000 | 190000 | 5001 | CN2 | CN2 | CN2 | CN2 | CN0 | CN2 | CN2 | CN2 | CN2 | CN2 | CN2 | CN2 | CN2 |
| 54 | mg35 | 131000 | 134000 | 3001 | CN2 | CN3 | CN2 | CN2 | CN2 | CN2 | CN2 | CN2 | CN2 | CN2 | CN2 | CN2 | CN2 |
| 55 | mg35 | 174000 | 178000 | 4001 | CN2 | CN2 | CN2 | CN1 | CN0 | CN2 | CN2 | CN2 | CN2 | CN2 | CN2 | CN2 | CN2 |
| 56 | mg37 | 0 | 13000 | 13001 | CN2 | CN2 | CN2 | CN2 | CN2 | CN2 | CN2 | CN1 | CN2 | CN2 | CN2 | CN2 | CN2 |
| 57 | mg38 | 101000 | 105000 | 4001 | CN3 | CN3 | CN2 | CN2 | CN2 | CN2 | CN2 | CN2 | CN2 | CN2 | CN2 | CN2 | CN2 |
| 58 | mg38 | 69000 | 72000 | 3001 | CN2 | CN2 | CN2 | CN2 | CN2 | CN2 | CN0 | CN2 | CN2 | CN2 | CN2 | CN2 | CN2 |
| 59 | mg38 | 167000 | 172000 | 5001 | CN2 | CN2 | CN2 | CN2 | CN2 | CN2 | CN1 | CN2 | CN2 | CN2 | CN2 | CN2 | CN2 |
| 60 | mg39 | 209000 | 214000 | 5001 | CN2 | CN2 | CN2 | CN2 | CN2 | CN2 | CN2 | CN2 | CN2 | CN2 | CN3 | CN3 | CN2 |
| 61 | mg39 | 216000 | 219555 | 3556 | CN2 | CN2 | CN2 | CN2 | CN2 | CN2 | CN2 | CN2 | CN2 | CN2 | CN3 | CN3 | CN2 |
| 62 | mg40 | 165000 | 169000 | 4001 | CN5 | CN2 | CN2 | CN2 | CN2 | CN2 | CN2 | CN2 | CN2 | CN2 | CN1 | CN2 | CN2 |
| 63 | mg40 | 129000 | 132000 | 3001 | CN1 | CN2 | CN2 | CN2 | CN2 | CN2 | CN2 | CN2 | CN2 | CN2 | CN2 | CN2 | CN2 |
| 64 | mg41 | 0 | 16000 | 16001 | CN2 | CN2 | CN2 | CN2 | CN2 | CN4 | CN2 | CN2 | CN2 | CN2 | CN2 | CN2 | CN2 |
| 65 | mg45 | 147000 | 150000 | 3001 | CN2 | CN2 | CN2 | CN2 | CN3 | CN2 | CN2 | CN2 | CN2 | CN2 | CN2 | CN2 | CN2 |
| 66 | mg46 | 72000 | 75000 | 3001 | CN2 | CN2 | CN2 | CN2 | CN1 | CN2 | CN2 | CN2 | CN2 | CN2 | CN2 | CN2 | CN2 |
| 67 | mg51 | 154000 | 162000 | 8001 | CN2 | CN2 | CN2 | CN2 | CN2 | CN3 | CN2 | CN2 | CN2 | CN2 | CN1 | CN2 | CN2 |
| 68 | mg52 | 138000 | 163942 | 25943 | CN2 | CN2 | CN2 | CN2 | CN1 | CN2 | CN2 | CN2 | CN2 | CN2 | CN2 | CN2 | CN2 |
| 69 | mg53 | 128000 | 131000 | 3001 | CN1 | CN2 | CN2 | CN2 | CN2 | CN2 | CN2 | CN2 | CN2 | CN2 | CN2 | CN2 | CN2 |
| 70 | mg53 | 133000 | 139000 | 6001 | CN1 | CN2 | CN2 | CN2 | CN2 | CN2 | CN2 | CN2 | CN2 | CN2 | CN2 | CN2 | CN2 |
| 71 | mg53 | 7000 | 11000 | 4001 | CN1 | CN0 | CN2 | CN1 | CN4 | CN6 | CN2 | CN2 | CN0 | CN2 | CN2 | CN2 | CN2 |
| 72 | mg58 | 0 | 10000 | 10001 | CN2 | CN1 | CN3 | CN1 | CN0 | CN3 | CN2 | CN2 | CN1 | CN2 | CN3 | CN3 | CN2 |
| 73 | mg61 | 120000 | 127393 | 7394 | CN1 | CN2 | CN2 | CN2 | CN2 | CN2 | CN2 | CN2 | CN0 | CN2 | CN2 | CN2 | CN2 |
| 74 | mg67 | 105000 | 110095 | 5096 | CN2 | CN2 | CN2 | CN2 | CN2 | CN3 | CN2 | CN2 | CN2 | CN2 | CN2 | CN2 | CN2 |
| 75 | mg68 | 103000 | 107941 | 4942 | CN2 | CN2 | CN2 | CN2 | CN2 | CN3 | CN2 | CN2 | CN2 | CN2 | CN3 | CN2 | CN2 |
| 76 | mg71 | 28000 | 31000 | 3001 | CN2 | CN2 | CN2 | CN2 | CN0 | CN2 | CN2 | CN2 | CN0 | CN2 | CN2 | CN2 | CN2 |
| 77 | mg71 | 0 | 20000 | 20001 | CN2 | CN2 | CN2 | CN2 | CN0 | CN2 | CN2 | CN2 | CN0 | CN2 | CN2 | CN2 | CN2 |
| 78 | mg72 | 7000 | 11000 | 4001 | CN2 | CN2 | CN2 | CN2 | CN0 | CN2 | CN2 | CN2 | CN2 | CN2 | CN2 | CN2 | CN2 |
| 79 | mg75 | 31000 | 60957 | 29958 | CN2 | CN2 | CN2 | CN2 | CN2 | CN2 | CN2 | CN2 | CN2 | CN2 | CN2 | CN2 | CN0 |
| 80 | mg76 | 43000 | 46000 | 3001 | CN2 | CN2 | CN2 | CN2 | CN0 | CN2 | CN2 | CN2 | CN2 | CN2 | CN2 | CN2 | CN2 |
| 81 | mg76 | 53000 | 59101 | 6102 | CN2 | CN2 | CN2 | CN2 | CN0 | CN2 | CN2 | CN2 | CN2 | CN2 | CN2 | CN2 | CN2 |
| 82 | mg77 | 42000 | 48000 | 6001 | CN2 | CN2 | CN2 | CN2 | CN2 | CN3 | CN2 | CN2 | CN2 | CN2 | CN2 | CN2 | CN2 |
| 83 | mg77 | 9000 | 23000 | 14001 | CN0 | CN2 | CN2 | CN2 | CN2 | CN2 | CN2 | CN2 | CN2 | CN2 | CN2 | CN2 | CN2 |
| 84 | mg78 | 0 | 11000 | 11001 | CN2 | CN2 | CN3 | CN2 | CN2 | CN2 | CN6 | CN2 | CN6 | CN2 | CN6 | CN6 | CN2 |
| 85 | mg79 | 0 | 11000 | 11001 | CN2 | CN2 | CN2 | CN2 | CN2 | CN2 | CN2 | CN2 | CN4 | CN2 | CN2 | CN2 | CN2 |
| 86 | mg85 | 23000 | 30000 | 7001 | CN2 | CN2 | CN2 | CN2 | CN1 | CN2 | CN2 | CN2 | CN1 | CN2 | CN2 | CN2 | CN2 |
| 87 | mg85 | 12000 | 16000 | 4001 | CN2 | CN2 | CN2 | CN2 | CN2 | CN2 | CN2 | CN2 | CN1 | CN2 | CN2 | CN2 | CN2 |
| 88 | mg86 | 22000 | 26000 | 4001 | CN2 | CN2 | CN2 | CN2 | CN2 | CN3 | CN2 | CN2 | CN2 | CN2 | CN2 | CN2 | CN2 |
| 89 | mg86 | 26000 | 29728 | 3729 | CN2 | CN2 | CN2 | CN2 | CN2 | CN3 | CN2 | CN2 | CN2 | CN2 | CN2 | CN2 | CN2 |
| 90 | mg87 | 0 | 24367 | 24368 | CN1 | CN2 | CN3 | CN0 | CN0 | CN2 | CN3 | CN1 | CN0 | CN2 | CN3 | CN3 | CN0 |
| 91 | mg92 | 0 | 13200 | 13201 | CN2 | CN2 | CN2 | CN2 | CN1 | CN2 | CN2 | CN2 | CN2 | CN2 | CN2 | CN2 | CN2 |
| 92 | mg195 | 0 | 5600 | 5601 | CN2 | CN2 | CN2 | CN2 | CN2 | CN2 | CN2 | CN2 | CN2 | CN2 | CN2 | CN2 | CN0 |
